# How resource abundance and stochasticity affect organisms’ range sizes

**DOI:** 10.1101/2023.11.03.565563

**Authors:** Stefano Mezzini, Chris H. Fleming, E. Patrícia Medici, Michael J. Noonan

**Author notes:** Correspondence: Michael J. Noonan < >.

## Abstract

The amount of space organisms use is thought to be tightly linked to the availability of resources within their habitats, such that organisms living in productive habitats generally require less space than those in resource-poor habitats. This hypothesis has widespread empirical support, but existing studies have focused primarily on responses to the mean amount of resources, while responses to the variance around the mean are still largely unknown. This is not a trivial oversight. Organisms adjust to variable environmental conditions, so failing to consider the effects of resource (un)predictability can result in a limited understanding of an organism’s range size, which challenges ecological theory and applied conservation alike. In this study, we leverage the available literature to provide a unifying framework and hypotheses for the effect of mean and variance in resources on range sizes. Next, we use simulated movement data to demonstrate how the combined effects of mean and variance in resource abundance interact to shape predictable patterns in range size. Finally, we use real-world tracking data on a lowland tapir (*Tapirus terrestris*) from the Brazilian Cerrado to show how this framework can be applied to better understand the movement ecology of free-ranging animals.

## Introduction

The amount of resources an organism is able to access is a strong determinant of its odds of survival and reproduction. Resource limitations can cause individuals to experience a negative energetic balance, which can then result in lower fitness [1,2], altered physiology [2–5], lower chance of reproduction [2,6–8], and even death [9,10]. Thus, many organisms adapt their behaviors and/or physiology in response to changes in local resource abundance to ensure their needs are met.

While there are many ways that individuals can respond to resource availability, move-ment represents one of the most readily available traits that species can adjust [11–13]. The relationship between organisms’ movement and resource abundance has long been of inter-est to biologists. In his seminal paper, [14] considered the search for food as the primary driver for movement within an organism’s home range. Three decades after, [15] suggested that change in resource abundance drives how organisms decide where to live and when to reproduce. Two years later, [16] proposed that the simplest relationship between resource abundance and an organism’s home-range size is

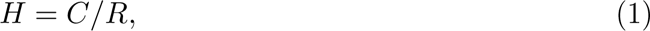

where *H* is the organism’s home-range size, *C* is the organism’s resource consumption (kcal day^−1^), and *R* is the resources the organism can access (kcal day^−1^ unit area^−1^). Harestad and Bunnel’s model is simple to conceptualize, and it allows for testable predictions, but few studies are structured around a set of theoretical expectations such as Harestad and Bunnel’s hypothesis. Many researchers have since demonstrated that organisms adapt their range sizes in response to resources abundance, but results are typically reported as independent, novel findings. Perhaps more problematic is the fact that, while much work has been done on estimating organisms’ responses to mean resource abundance, there is little information on how organisms respond to variance around the mean [i.e., resource stochasticity, but see: 17,18,19]. Thus, there remains a need for clear hypotheses of the effects of both resource abundance and stochasticity on organisms’ range sizes.

Here, we refer to a location’s average amount of resources as “resource abundance”, while we use the phrase “resource stochasticity” to indicate the variability in resources after accounting for changes in the mean. We argue that, on its own, a habitat’s resource abundance is not sufficient to assess the habitat’s quality, nor make predictions about how much space an organism might use. To see this, consider, for instance, a herbivore grazing in a grassland with relatively low but constant forage availability (i.e., low mean and variance). This individual will adopt different behaviors and adaptations if it lived in a desert with equally scarce forage but rare, sudden, and strong pulses of resources (i.e., low mean and high stochasticity). In the grassland, the grazer may require a large but constant home range size as it moves between patches in search of food, while in the desert it may switch between dispersal in search for high-resource patches and short-term range residency within patches [*sensu* 12,see 20,21,22]. Previous studies suggest that resource stochasticity may decrease organisms’ fitness and landscapes’ energetic balances [e.g., 23], but there is still limited empirical evidence to support this hypothesis [19,but see: 24,25].

In this paper, we illustrate how an organism’s range size can be expected to depend on both the abundance and unpredictability of resources. First, we set the theoretical background necessary for the successive sections by introducing key concepts and notation. Next, we provide a review of the effects of resource abundance on range sizes while suggesting a simple and unifying hypothesis. Afterwards, we provide a review of the effects of resource stochasticity on organisms’ range sizes while suggesting a second simple and unifying hypothesis. Subsequently, we support the two hypotheses using quantitative, simulated responses in range size to changes in resource abundance and stochasticity. Finally, we demonstrate how this framework can be used in practice to describe the movement ecology of a lowland tapir (*Tapirus terrestris*) from the Brazilian Cerrado [26].

### Resources as a random variable

Resources are often unpredictable (and difficult to quantify), since they depend on various factors which cannot be accounted for easily, including climate [7,27,28], weather [28,29], competitive pressure [30,31], and differences in energetics at among individuals [7] and species [32]. Thus, we can treat the amount of resources *R* at a given point in time (*t*) and space (location vector *u⃗*) as a random variable, denoted as *R*(*t*, *u⃗*). Treating resources as a random variable allows us to leverage techniques from probability theory and statistics, such as the expectation of a random variable (i.e., its mean) and its variance around the mean. We indicate the expected value and variance of random variable *R* using E(*R*) and Var(*R*), respectively, and we use *µ*(*t*, *u⃗*) and *σ*^2^(*t*, *u⃗*) to indicate them as functions of time (*t*) and space (*u⃗*). Appendix A defines and expands on the concepts of probability distributions, expected value, variance, and provides examples of them for gamma and beta distributions.

### Effects of resource abundance, E(*R*)

While organisms’ needs vary greatly between taxonomic groups, some needs are essential for the growth, survival, and reproduction of most organisms. All heterotrophic organisms require sources of chemical energy (i.e., food), water, and various limiting nutrients [33–35]. As the abundance of essential resources fluctuates, motile organisms can move to new locations or ‘patches’ to meet their requirements [12,36], but they must also account for costs of movement [37].

Fig. 1 illustrates our first of two hypotheses, which is similar to that presented by [16]. When E(*R*) is high, we expect organisms’ ranges to be relatively small and near the smallest amount of space required to survive [24,25,e.g., 38]. Like [16], we also expect organisms’ range sizes to increase nonlinearly as E(*R*) decreases, but we highlight that organisms may adopt different behaviors at low values of E(*R*). These behaviors include maximal home range expansion [30,home range size is limited by vagility, habitat structure, competition, and predation, e.g., 31,39,40], migration [41–43], and nomadism [20,22,44,45]. It is unclear when organisms switch from range residency to migration or nomadism (or vice-versa), but understanding the gradient among these types of movement is necessary for quantifying the effect of resource abundance on organisms’ range size and movement behavior [mammals: 46,moose, *Alces alces*: 20,eagles, *Haliaeetus leucocephalus*: 21,47,lesser flamingos, *Phoeniconaias minor*: 48]. Still, switches from range residency to nomadism (or vice-versa) will occur over evolutionary timescales rather than over an organism’s lifespan (Fig. 1), since larger ranges require greater vagility, which, in turn, is facilitated by the development of morphological features such as hinged joints and elongated limbs [32,49–51].

**Figure 1:**
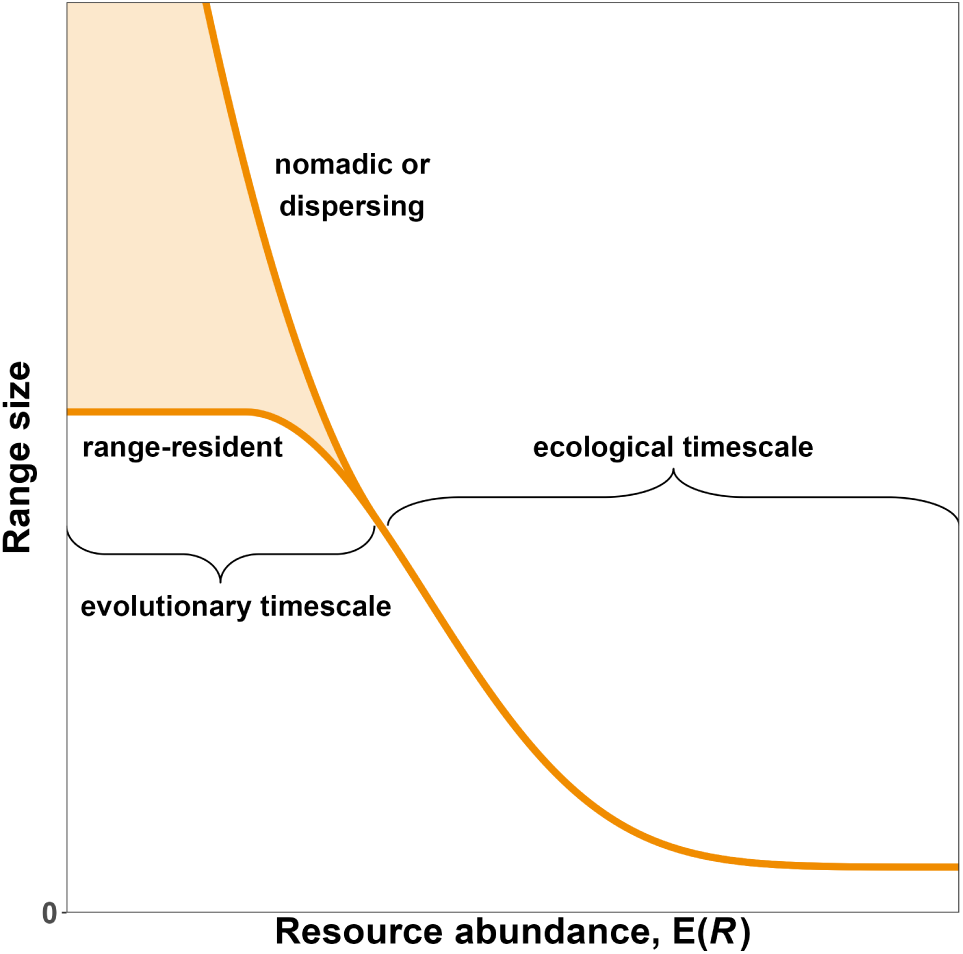
Hypothesized range size of an organism as a function of resource abundance, E(*R*). We expect low values of E(*R*) to result in a large range, since organisms are forced to explore large areas to collect the resources they require to survive, whether they be range-resident, nomadic, or migratory. As E(*R*) increases, range size should decrease nonlinearly until it reaches the minimum amount of space required by the organism to survive. Note that the relationship between E(*R*) and range size cannot be linear because it would require range size to be negative for high values of E(*R*).

Overall, the hypothesis that range size decreases with resource abundance, E(*R*), is com-monly accepted and well supported, but many studies assume a linear relationship [19,e.g., 38,52–54]. This is problematic because, conceptually, the relationship between range size and E(*R*) must be nonlinear, since: (1) there is an upper limit to how much space an organ-ism is able to explore in its finite lifetime and (2) the minimum amount of space it requires to survive is necessarily greater than zero [24,25,see: 55,56,57, and contrast them to the estimates based on linear models listed above]. Consequently, we suggest analysts use models that account for this nonlinearity when estimating the effects of resource abundance on range size.

### Effects of resource stochasticity, Var(*R*)

Assuming resource stochasticity is constant over time and space can be a useful simplification of relatively stable environments or when information on how E(*R*) changes is limited and estimating changes in Var(*R*) is unreasonable. However, such an assumption is likely not realistic, since Var(*R*) often differ across space and over time. Generally, bounded qualities quantities have correlated means and variances, as in the case of random variables that are strictly positive (e.g., Gamma and Poisson) or fully bounded (e.g., beta). See Appendix A for more information.

Recognizing changes in Var(*R*) helps account for the residual, fine-scale variation in *R* after accounting for trends in the large-scale average *R* [e.g., variations in plant phenology between years after accounting for mean seasonal trends, see 58]. However, when both E(*R*) and Var(*R*) change over time (fig. A2), disentangling changes in E(*R*) and Var(*R*) is not simple [59]. Statistically, this confound occurs because the more change one attributes to *µ*(*t*, *u⃗*) (i.e., the wigglier it is), the smaller *σ*^2^(*t*, *u⃗*) becomes. Conversely, the smoother *µ*(*t*, *u⃗*) is, the larger *σ*^2^(*t*, *u⃗*) becomes. Biologically, it is important because an organism’s perception scale determines whether it attributes a change in *R* to a trend in E(*R*) or as a stochastic event [i.e., due to Var(*R*); see [58]]. An organism’s perception of changes in *R* will also depend strongly on the its cognitive capacities and memory [9,60–63]. Whether an organism is able to predict trends in *σ*^2^(*t*, *u⃗*) or not, environmental variability is thought to reduce a landscape’s energetic balance [23], which, in turn, decreases organisms’ fitness [e.g., 10] and increases their range size. While this behavioral response occurs with both predictable and unpredictable stochasticity, extreme and rare events are more likely to have a stronger effect due to their unpredictability and magnitude [64,65]. A few recent studies support these hypotheses [23,28,45,66], but many of them are limited in geographic and taxonomic scales, so the extent to which these preliminary findings can be generalized is currently unknown. Thus, there remains a need for developing a more complete understanding of how organisms’ range sizes changes with environmental stochasticity.

Similarly to E(*R*), we hypothesize Var(*R*) has a nonlinear effect on an organism’s range size. When Var(*R*) is low enough that *R* is relatively predictable, we expect organisms to be range-resident with small home ranges, and we do not expect small changes in Var(*R*) to have a noticeable effect. As resources become increasingly unpredictable, we expect home range size to increase progressively faster (fig. 2) because: (1) as Var(*R*) increases, the chances of finding low *R* increase superlinearly, (2) the added movement required to search for food increases organisms’ energetic requirements, and (3) stochasticity reduces an organism’s ability to specialize and reduce competition for *R* [67]. If resources remain highly unpredictable over long periods of time (e.g., multiple lifespans), organisms may evolve or develop new and consistent behaviors (e.g, nomadism) or adaptations (e.g., increased fat storage or food caching) to buffer themselves against times of unpredictably low *R*. Conversely, if changes in *σ*^2^(*t*, *u⃗*) are sufficiently predictable, organisms may learn to anticipate and prepare for times of greater stochasticity by pre-preemptively caching food, reducing energetic needs, migrating, or relying on alternative food sources [e.g., 68].

**Figure 2:**
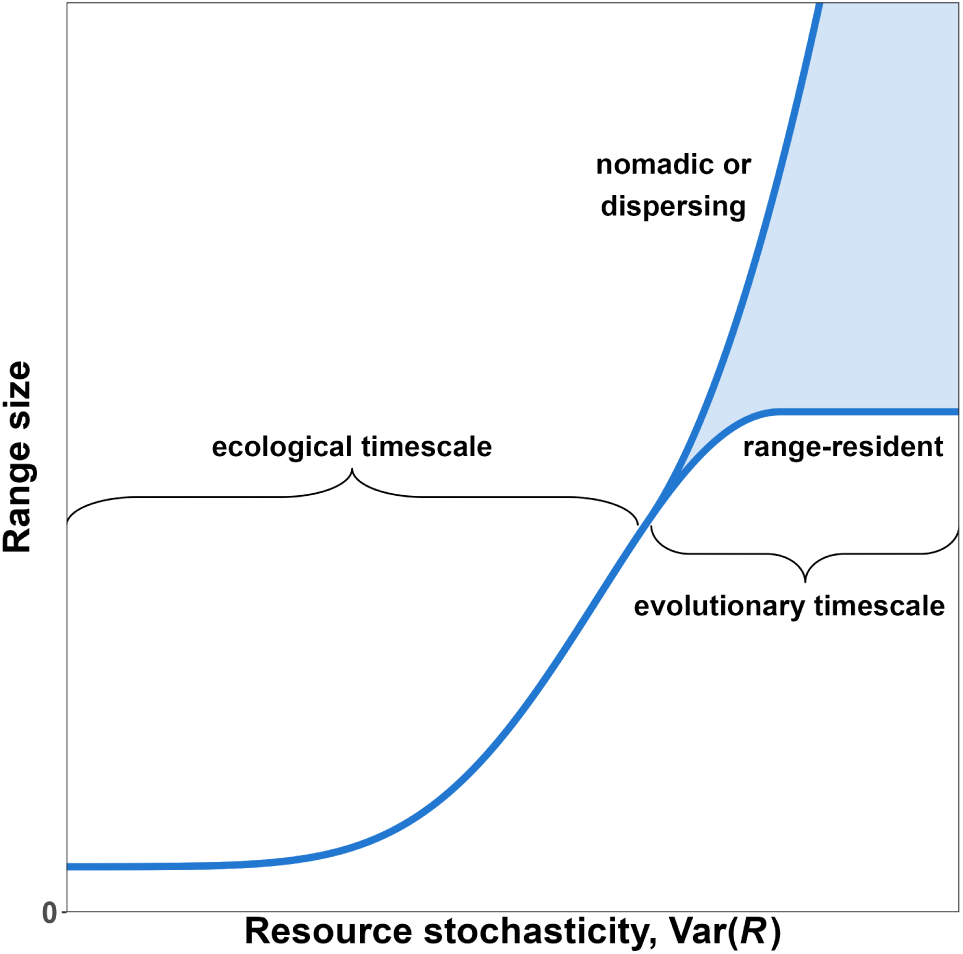
Hypothesized range size of an organism as a function of resource stochasticity, Var(*R*). We expect low values of Var(*R*) to result in small home-rages, since organisms are able to depend on relatively predictable resources. As Var(*R*) increases, range size should increase nonlinearly, whether this results in an expansion of the home range (in the case of range-resident organisms) or a switch to a larger range via dispersal, nomadism, or migration. Note that the relationship between Var(*R*) and range size cannot be linear because it would require range size to be negative for low values of Var(*R*).

### Interactive effects of E(*R*) and Var(*R*)

We have provided the case for why both E(*R*) and Var(*R*) should be expected to affect organisms’ range size, but we presented the two parameters as independent drivers of move-ment. However, organisms may respond to changes in *σ*^2^(*t*, *u⃗*) more when resources are scarce than when they are abundant. Consequently, an organism’s movement behavior is likely to be a function of not only the marginal effects of E(*R*) and Var(*R*) but also their interactive effects. A highly unpredictable habitat may be very inhospitable if resources are poor, but Var(*R*) may have little effect if resources are stochastic but always abundant. Thus, we expect Var(*R*) to have a stronger effect on range size when E(*R*) is low, and less of an effect when E(*R*) is high. We explore this interaction effect more in the following section.

### Simulating responses to E(*R*) and Var(*R*)

To support our hypotheses of how organisms’ range sizes are affected by E(*R*), Var(*R*), and the interaction effect of E(*R*) and Var(*R*), we present the results from a series of quantitative simulations. To start, we used the ctmm package [69] for R [70] to generate 200 tracks (see Appendix B for sensitivity analyses) from an Integrated Ornstein-Uhlenbeck movement model [IOU model, see 71]. The IOU model’s correlated velocity produced realistic tracks with directional persistence, but, unlike Ornstein-Uhlenbeck (OU) and Ornstein-Uhlenbeck Foraging (OUF) models, IOU models do not produce spatially stationary movement, so the organism is not to range-resident. Consequently, each track is spatially unrestricted and can be interpreted as purely exploratory or memoryless movement.

Each of the 200 tracks were placed on a grid with common starting point ⟨0, 0⟩ (fig. B1). Each time the simulated individual moved to a new cell, it collected *R* resources sampled from a Gamma distribution. The mean and variance of the distribution were defined by a series of deterministic functions *µ*(*t*) and *σ*^2^(*t*) (orange and blue lines in fig. 3). The value of *t* was constant within each set of 200 tracks, so the distribution *R* was sampled from was independent of both the organism’s location and its time spent moving. Tracks were truncated once the organism reached satiety, and the organism was given enough time to return to ⟨0, 0⟩ independently from the following track (section 2.1 of Appendix B). Finally, we fit an OUF movement model [72] to the set of tracks to calculate the 95% Gaussian home-range size using the formula

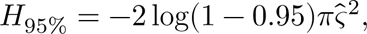

**Figure 3:**
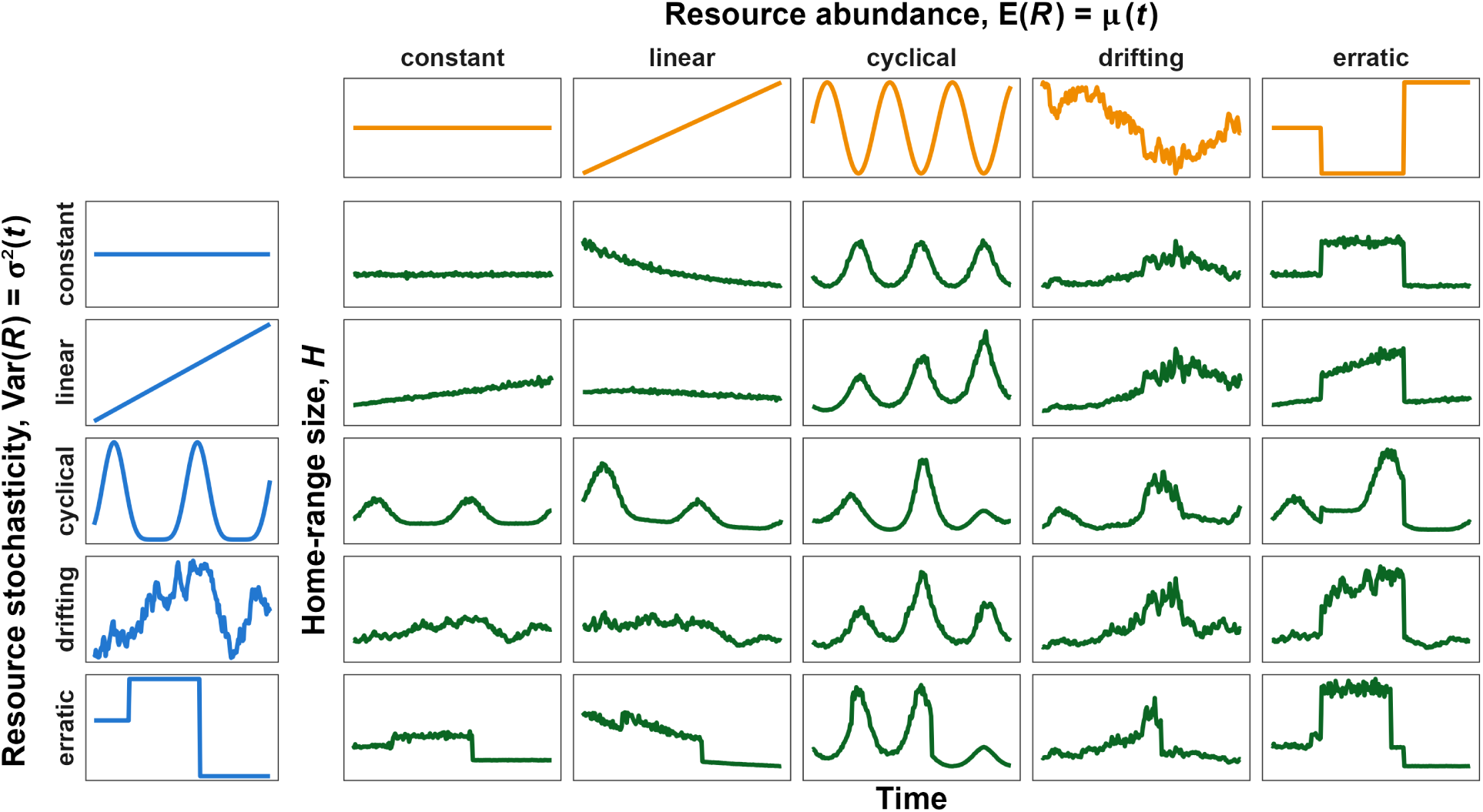
Simulated home-range sizes, *H*, of an organism living in habitats where the mean and variance in resources are constant, linearly increasing, cyclical, drifting, or erratic over time (but homogenous over space for a given *t*). Note how *H* decreases nonlinearly as *µ*(*t*) increases and increases nonlinearly as *σ*^2^(*t*) increases. Additionally, the variance in *H* is higher when *µ*(*t*) is lower or *σ*^2^(*t*) is higher, and changes in *σ*^2^(*t*) have greater impacts when *µ*(*t*) is low.

where 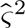 is the positional variance estimated by the movement model.

We designed the simulations to estimate the effects of E(*R*) and Var(*R*) in simplistic environments where organisms could only respond by searching for longer periods of time. Consequently, we made the following assumptions:

1) Environments are homogeneous for a given *t*. Given *t*, E(*R*) = *µ*(*t*) and Var(*R*) = *σ*^2^(*t*) are constant over space and within each set of 200 tracks, but *R* is random and follows a Γ(*µ*(*t*), *σ*^2^(*t*)) distribution.
2) The are no external pressures on the simulated organism. Resources do not deplete, and there is no competition nor predator avoidance.
3) The organism has a fixed daily energetic requirement that is independent of movement rates, and it cannot alter its metabolism or physiology. Additionally, the organism does not have energetic reserves, so excess resources cannot be carried over to the next track or *t*.
4) The organism is range-resident and can only respond to changes in E(*R*) and Var(*R*) by altering its home-range size. The organism does not disperse or abandon a range.
5) The organism’s movement is simplistic. The organism’s movement speed and direction are stochastic and independent of E(*R*) and Var(*R*).
6) The organism has no perceptive range or memory. It is unable to detect, learn, or predict where resources are abundant (high E(*R*)) or reliable (low Var(*R*)) over time or space.
7) Animals only move to search for food or return to the center of their home-range after reaching satiety.

Additional information is provided in Appendix B, including the directed acyclical graph [see fig. B6 and 73] we used to infer causal the mechanisms of changes in *H* and estimate the direct effects of E(*R*) and Var(*R*) on *H* (contrast the graph with fig. C3 and the empirical case study below).

Fig. 3 shows how simulated home-range size, *H*, responded to changes in *µ*(*t*) and *σ*^2^(*t*) in scenarios where both functions can remain constant, increase linearly, oscillate cyclically, drift stochastically, or change erratically. The top row (constant Var(*R*)) shows how *H* varies for different trends in *µ*(*t*) while Var(*R*) remains constant (like in fig. A1). As E(*R*) increases at a constant slope (linear *µ*(*t*)), *H* decreases nonlinearly, with larger changes when E(*R*) is low, until it approaches the minimum size required by the organism. Also note how the noise in the green lines also decreases as E(*R*) increases.

The leftmost column of fig. 3 (constant E(*R*)) illustrates the effects of Var(*R*) on *H* while E(*R*) remains constant. Overall, both mean *H* and the variance around it increase with *σ*^2^(*t*) (most visible with constant E(*R*) and linear Var(*R*)). Similarly to resource-poor periods, times of greater stochasticity require the organism to move over larger areas for longer periods of time. Additionally, the greater in uncertainty in how much time and space the organism will require to reach satiety, or indeed whether an organism living in highly stochastic environments can even reach satiety within a finite amount of time.

The remaining panels in fig. 3 illustrate how E(*R*) and Var(*R*) jointly affect *H* and how confusing the effects can be. Since E(*R*) and Var(*R*) have opposite effects on *H*, disentangling the effects can be particularly difficult when both parameters change in a correlated manner (e.g., linear E(*R*) and Var(*R*)). When both E(*R*) and Var(*R*) increase linearly, *H* initially increases since the effect of Var(*R*) is stronger, but then decreases as the effect of E(*R*) begins to dominate. Difficulties in disentangling the two effects are explored in greater depth in the case study in the following section.

Although the temporal trends in fig. 3 are complex and the effects of E(*R*) and Var(*R*) can be hard to disentangle, two simple relationships emerge when *H* is shown as a function of either E(*R*) or Var(*R*), rather than time (panels A and B of fig. 4). The estimated relationships follow the hypotheses we presented in figs. 1 and 2, although we found that the effect of Var(*R*) at average E(*R*) was linear with a slight sublinear saturation at high values of Var(*R*). However, notice that the effect of Var(*R*) on *E*(*H*) depends strongly on E(*R*) (panel C): When E(*R*) is low, E(*H*) is high and Var(*R*) does not have a strong effect, but when E(*R*) is high the effect of Var(*R*) on E(*H*) is exponential. Similarly, E(*H*) decreases exponentially with E(*R*) except when Var(*R*) is very high.

**Figure 4:**
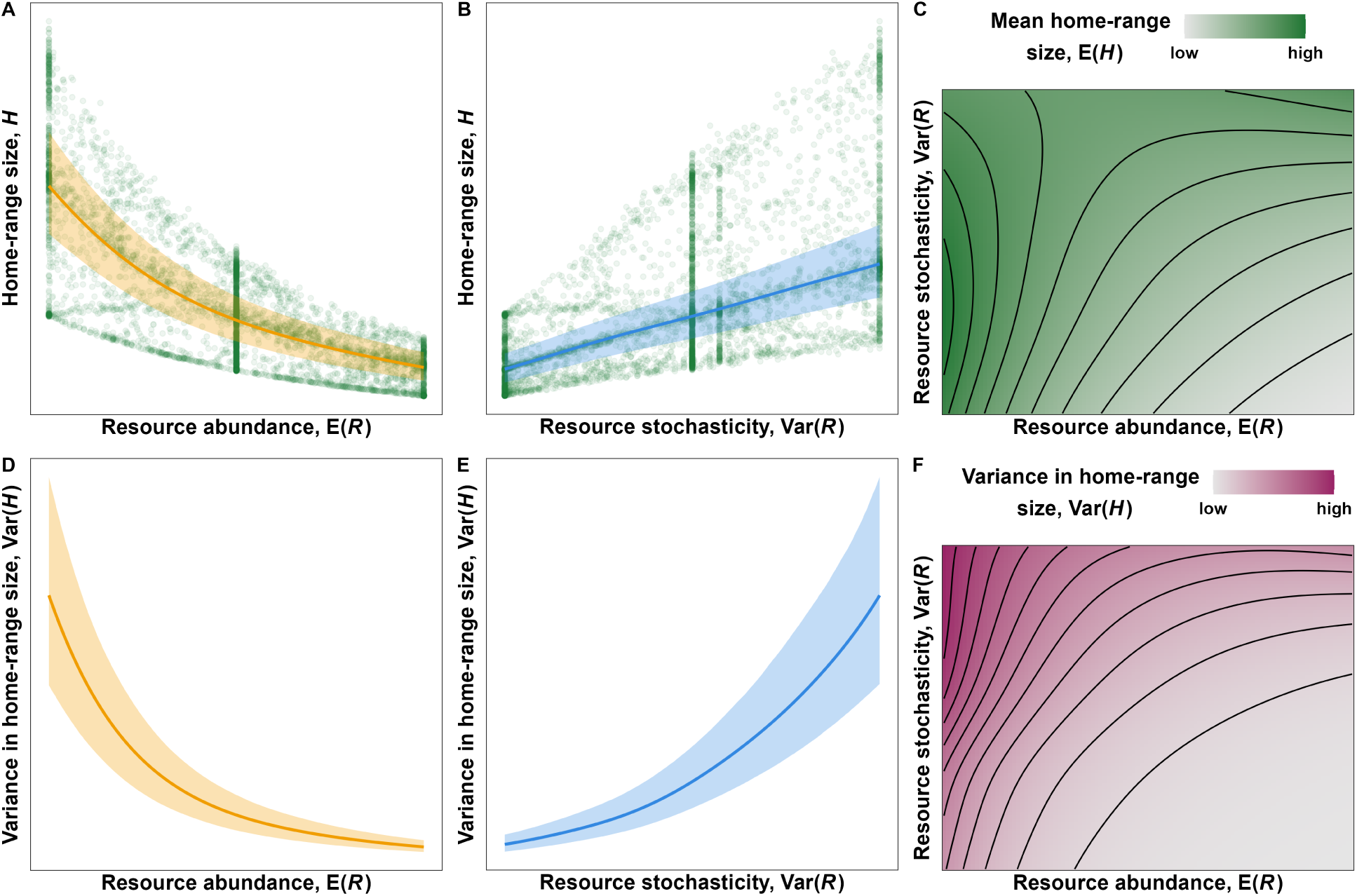
Effects of E(*R*) and Var(*R*) on on the mean (A-C) and variance (D-F) in simulated home-range size with 95% Bayesian credible intervals. While the estimated marginal effect of Var(*R*) on E(*H*) is sublinear (panel B), the effect of Var(*R*) is superlinear for high values of E(*R*) (panel C). The relationships were estimated using a Generalized Additive Model for Location and Scale with a Gamma location-scale family of distributions (mgcv∶∶gammals). Credible intervals were calculated using 10,000 samples from the posterior distribution while assuming multivariate Gaussian coefficients. Additional details on the model structure are provided in Appendix B.

As expected by the changes in the spread of the points in panels A and B of fig. 4, the variance in *H*, Var(*H*), also depends on E(*R*) and Var(*R*) (fig. 4D-F). Since we modeled *H* using a Gamma family of distributions, we expected Var(*H*) to increase with E(*H*), but the location-scale model removes the assumption of a constant mean-variance relationship (i.e., constant coefficient of variation, <u>^*µ*(*t*)^</u>. This allowed us to show that the effect of *R* on Var(*H*) is much stronger than the effect of *R* on E(*H*). Consequences of these effects are explored in the discussion section.

### A case study on a lowland tapir in the Brazilian Cerrado

The simulations in the section above support the hypotheses we presented in the introduction, but they are based on assumptions that are often not met in real natural environments. Organisms live in spatiotemporally heterogeneous and dynamic environments that promote the use of perceptual ranges, navigation, and memory. Together, these abilities result in selective space use that depends on resource availability [11] and resource depletion [12].

In this section, we test the hypotheses using empirical tracking data on a lowland tapir from the Brazilian Cerrado along with empirical estimates of E(*R*) and Var(*R*). We measure *R* using Normalized Difference Vegetation Index [NDVI, see 74], a remote-sensed measure of landscape greenness, as a proxy for forage abundance. Appendix C contains additional information on how we modeled NDVI and the tapir’s movement using continuous-time movement models [69,75] and autocorrelated kernel density estimation [76–78].

Fig. 5 illustrates how a tapir in the Brazilian Cerrado adapts its 7-day home-range size to spatiotemporal changes in *µ*(*t*, *u⃗*) and *σ*^2^(*t*, *u⃗*) [telemetry data from the individual labelled as “Anna” in the dataset from 26]. Panels A and B show the changes in seven-day average mean and variance in NDVI, respectively, experienced by the tapir during the tracking period. The mean and variance in NDVI were estimated using a Generalized Additive Model for Location and Scale [GAMLS, 79] with a Beta family of distributions (NDVI values ranged from 0.3534 to 0.9475). Panel C shows the changes in the tapir’s 7-day home range over time. Note how the tapir uses more space during periods of lower NDVI (e.g., August 2017) and less space during periods with high NDVI (January 2018). Additionally, when resources are scarce and highly unpredictable (August 2018), the tapir uses up to 5 times more space than when resources are abundant and predictable (e.g., January 2018). Finally, panels D and E show the estimated (marginal) effects of *µ*(*t*, *u⃗*) and *σ*^2^(*t*, *u⃗*) on the tapir’s 7-day home-range size. Since *µ*(*t*, *u⃗*) and *σ*^2^(*t*, *u⃗*) are correlated (panel F) and spatiotemporally autocorrelated (panels A, B, and F), the effects of *R* on *H* should be modeled carefully. To avoid over-fitting the model, we constrained the smooth effects of *µ*(*t*, *u⃗*) and *σ*^2^(*t*, *u⃗*) and their interaction effect to a small basis size (k = 3). Additional information is provided in appendix C. The results presented in panels D-F of fig. 5 match our findings from the simulations: The tapir’s 7-day home range decreases with *µ*(*t*, *u⃗*) and increases with *σ*^2^(*t*, *u⃗*), and the effect of *µ*(*t*, *u⃗*) depends on *σ*^2^(*t*, *u⃗*), and vice-versa. Alone, *µ*(*t*, *u⃗*) and *σ*^2^(*t*, *u⃗*) cause the tapir to double her home range (panels D and E), but together they result in an approximate 15-fold change in home-range size (observed range: 0.8 to 12.4 km^2^; see panel F). Additionally, note how high NDVI values (> 0.8) cause *σ*^2^(*t*, *u⃗*) to have little to no effect on home-range size, as indicated by the vertical contour line in panel F.

**Figure 5:**
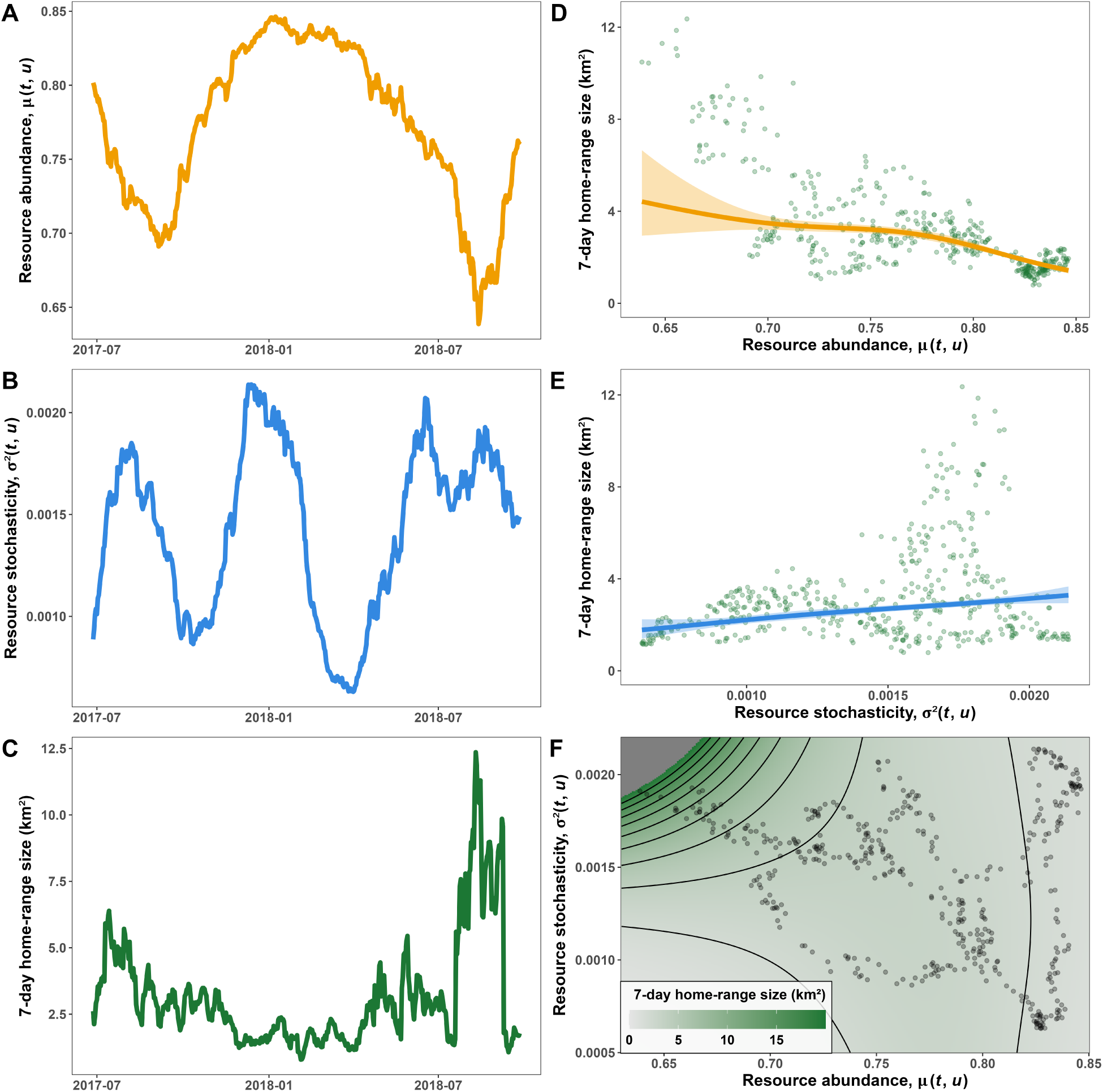
Effects of *µ*(*t*, *u⃗*) and *σ*^2^(*t*, *u⃗*) on the home-range size of a lowland tapir (*Tapirus terrestris*). (A) Trends in resource abundance over time, *µ*(*t*, *u⃗*), estimated as the average mean NDVI at the locations visited by the tapir during a seven-day period. (B) Varince in resources over time, *σ*^2^(*t*, *u⃗*), esimated as the average variance in NDVI at the locations visited by the tapir during a seven-day period. (C) Seven-day 95% home range estimated using Autocorrelated Kernel Density Estimation. (D, E) Estimated marginal effects of *µ*(*t*, *u⃗*) and *σ*^2^(*t*, *u⃗*) on home-range size. The model accounted for the marginal effects of *µ*(*t*, *u⃗*) and *σ*^2^(*t*, *u⃗*) and their interaction effect. (F) Estimated home-range size in response to changes in both *µ*(*t*, *u⃗*) and *σ*^2^(*t*, *u⃗*). Note how the effect of *σ*^2^(*t*, *u⃗*) is more pronounced when *µ*(*t*, *u⃗*) is low. See Appendix C for additional information. The tapir movement data corresponds to the individual named “Anna” from the Cerrado sample of Medici *et al.* (2022).

## Discussion

The amount of space organisms use is determined by a multitude of factors [13], but the search for resources is often a main driver of animal how much and where organisms move. This paper builds on earlier theoretical work [15,e.g., 16,17] and presents two hypotheses that describe the effects of resource abundance and stochasticity on organisms’ range sizes. We use quantitative simulations and an empirical case study to support the hypotheses and show that, together, they provide a simple framework for understanding how motile organisms adapt their movement in dynamic environments. Separately, resource abundance and stochasticity have simple but opposing effects on organisms’ range sizes: *H* decreases with E(*R*) and increases with Var(*R*). Together, the degree to which E(*R*) affects *H* depends on Var(*R*), and vice-versa, so organisms’ responses to resource dynamics can be complex. The simulated and empirical results suggest qualitatively similar marginal effects of E(*R*) and Var(*R*), but there are differences in the estimated interactive effects. In the simulated data, Var(*R*) has little effect when E(*R*) is low and a strong effect when E(*R*) is high, while the opposite is true for the empirical data. This difference is due to two reasons. Firstly, the shape and symmetry of bounded distributions such as Gamma (*R* > 0) and Beta (0 < *R* < 1) distributions depend on both E(*R*) and Var(*R*) (figs. A3, A4), but Var(*R*) does not affect the shape of a Gamma distribution as much if E(*R*) is low (fig. B3). Secondly, and perhaps more interestingly, the simulation approach does not account for real-world adaptations to E(*R*) and Var(*R*) such as selective space use, which we account for in the empirical approach. Below we discuss the strengths and limitations of each approach.

### Strengths and limitations of the simulation-based approach

Our simulations are based on a simplistic environment with many assumptions that allowed us to estimate how resource abundance and stochasticity affect organisms’ home-range sizes if organisms can only respond to changes by adapting the amount of time spent searching for food (with no energetic cost to movement). The use of continuous-time movement models coupled with few drivers of movement supported realistic data that could be explained by straightforward causal models. The absence of confounding variables (e.g., predator avoidance, territoriality, competition, landscape connectivity) or sample size limitation allowed us to ensure estimates were accurate and robust (sensitivity analysis available in Appendix B).

Deviations from the simulations offer a means of detecting when the underlying assumptions are inappropriate and how additional factors may affect organisms’ responses to changes in E(*R*) and Var(*R*). For example, energetic costs of movement are often non-negligible and depend on organism size [37], movement speed [37], and ambient temperature [1,80]. In addition, an organism may alter its movement behavior, physiology, and energetic needs to buffer itself against changes in E(*R*) and Var(*R*) by using space selectively [66,81–83] and adapting their behavior and physiology over time [15,67]. Before or during periods of scarcity, organisms may cache resources [84], build up fat reserves [42], enter states of dormancy [85–87], or even pause fetal growth [7]. However, organisms may be unable to respond to changes in E(*R*) and Var(*R*) optimally due to various reasons, including limited perceptive range [59], lack of experience [9,44,61–63,88], avoidance of competitors and predators [11,89], or a physiology that is not amenable to things like hibernation or fat storage. Thus organisms may relocate their range to a sub-optimal location [30,31,90,91], which may exacerbate the effects of E(*R*) and Var(*R*) on both mean range size and the variance around it.

### Strengths and limitations of the empirical approach

There are two main advantages of taking an empirical approach. Firstly, modeling real-world animal movement data can produce scale-appropriate and easily interpretable estimates. Secondly, empirical models directly quantify the effects of E(*R*), Var(*R*), and confounding variables without having to design complex and time-consuming simulations. However, it is not always possible to quantify confounding variables. For exaple, while there may be some appropriate proxies of competition, such as density of competitors, these variables may be hard to quantify, and they may not account for the confounding effects appropriately (i.e., the presence of competitors may not reflect competitive pressure). This is problematic if one is interested in estimating the direct causal effect of E(*R*) and Var(*R*), which requires removing any non-negligible confounding effects [73].

Similarly, if *R* is often non-measurable. Proxies of *R*, such as NDVI [74], which may introduce complexities. While *R* and NDVI are correlated for many species [42,43,88,e.g., 92,93,94], the relationship between the two can be weak [95], satellite-dependent [96], and nonlinear [96,97]. This complexity can introduce two sources of bias: ecosystem-level biases (indicated as *Z* in the directed acyclical graph in fig. C3) and satellite-level confounding variables (*S* in fig. C3). Examples of ecosystem-level biases are the effects of competition, predation, habitat connectivity, and movement costs, all of which can depend on habitat quality, and, consequently, be correlated nonlinearly to *R* and NDVI [32,98]. Resource-rich patches can attract larger amounts of competitors [11] and predators [18], which may, in turn, increase pressures from competition and predation [12,36]. However, such pressures may result in both an expansion of the range [32,98] or a contraction, since larger ranges can be harder to defend and result in higher movement costs [32,99] and encounter rates [100]. Satellite-level confounds include information loss due to coarse spatiotemporal resolution [96,97], satellite-level error [96,97,101], and other limitations of remote sensing (e.g., inability to quantify specific resources or small-scale resource depletion). However, nonlinear models such as Generalized Additive Models [102] can help account for preferences for intermediate values of remotely-sensed *R* [e.g., young grass rather than mature grasslands, see 96].

## Conclusion

The work presented here provides a unifying framework for viewing movement as a response to resource abundance and stochasticity. We provide realistic and flexible hypotheses of the effects of E(*R*) and Var(*R*) on organisms’ range sizes and movement behavior. We demonstrate that organisms’ range sizes decrease with resource abundance, increase with resource stochasticity, and that the effects of Var(*R*) can depend strongly on E(*R*).

Recent advances in computational power have greatly increased analysts’ ability to fit computationally demanding models [103,104] that allow biologists to move beyond only considering changes in mean conditions. By accounting for changes in stochasticity, we can start developing a more comprehensive understanding of how organisms adapt to the dynamic environments organisms live in, including recent changes in climate [105] and increases in the frequenct and intensity of extreme events [64,65,106–108].

## Supporting information

Appendix A: Concepts and definitions

Appendix B: Simulations

Appendix C: Empirical modeling

## Acknowledgements

We would like to thank Dr. Simon Wood for providing code to fit a Beta location-scale GAM despite not being involved directly with the project. Additionally, we thank all those who provided feedback on all posters, presentation, and writings related to this project. In particular, we thank all those who provided feedback on the manuscript and appendices despite not being authors, namely, in alphabetical order by first name: Aimee Chhen, Jessa Marley, Kim Hinz, Lauren Mills, Sarah Wyse, and Dr. Simon Wood. This work was supported by the NSERC Discovery Grant RGPIN-2021-02758 to MJN and funds from the University of British Columbia Okanagan, the Canadian Foundation for Innovation, Mitacs, and BC Parks.

## Code and data availability

All code and data used for this manuscript is available on GitHub at https://github.com/QuantitativeEcologyLab/hr-resource-stoch, with the exception of the tapir data, which is available at https://github.com/StefanoMezzini/tapirs.

