## Appendix A: Concepts and definitions for "How resource abundance and stochasticity affect organisms’ range sizes"

In this appendix, we presents the foundational concepts and terms used in the main manuscript. We introduce each concept without assuming a statistical or quantitative background to maximize accessibility and minimize misunderstandings and misinterpretations.

### Resources as a random variable

In statistics, random variables indicate random (i.e., unknown) quantities and are indicated with capital letters (e.g., $R$). Known values, such as realizations of random variables (i.e., known observations or instances), are indicated with lower-case letters (e.g., $r$). Using this notation, we can write the statement "the probability of random variable $R$ taking the value $r$" as $P(R = r)$.

In our simulations, we simulate $R$ using a Gamma distribution with time-dependent (but spatially homogeneous) mean $\mu(t)$ and variance $\sigma^2(t)$ (see the sections below), which we write as $R \sim \Gamma\Big(\mu(t), \sigma^2(t)\Big)$. Although Gamma distributions are more often parameterized using parameters shape and scale ($k > 0$ and $\theta > 0$) or shape and rate ($\alpha > 0$ and $\beta = 1/\theta > 0$), we use $\mathrm{E}(R) = k\theta$ and $\mathrm{Var}(R) = k\theta^2$ to facilitate visualizing the simulations within the context of resource abundance and stochasticity. However, note that $\mathrm{E}(R)$ and $\mathrm{Var}(R)$ are not independent because the variance depends strongly on the mean (and vice-versa). As the mean approaches zero from the positive side, the variance also does: $\mathrm{E}(R) \to 0^+ \iff k\theta \to 0^+ \iff k\theta^2 = 0^+ \iff \mathrm{Var}(R) = 0^+$. This assumption also holds biologically, since resources tend to be less variable when they are less abundant.

#### Probability distributions

Random variables are defined by specifying the distribution the variable is assumed to follow (e.g., Gaussian, Gamma, Poisson, Bernoulli). Since the variable is random, it can take multiple possible values, each with different probabilities. The set or range of values which have non-zero probabilities in a distribution is referred to as the distribution's support.

There are many distributions we can assign to $R$ depending on how we quantify it. For instance, if $R$ is the number of calories an animal is able to access from food in a given location, we can let $R$ follow a distribution with support positive real numbers, such as a Gamma or log-normal distribution. Alternatively, if we measure $R$ using the Normalized Difference Vegetation Index (NDVI, see Pettorelli et al. 2011), we should use a distribution with support over the interval $[-1, 1]$, since NDVI can only take on values between -1 and 1 (extremes included). However, there is no commonly used distribution with that support, so the best option is to rescale NDVI to (0, 1) and use a beta distribution (see the main text and Appendix C). If $R$ is a discrete integer variable, such as the number of prey in a location during a period of time, we can use a Poisson or negative binomial distribution.

**Expected resources, E($R$)**

Since the exact value of $R$ at a given time and location is unknown, comparing the magnitude in $R$ between two locations or time periods requires a quantitative measure of what values we believe $R$ will take, on average, in each of the two locations or time periods. The expectation of a random variable is the value one can expect the random variable to take, on average, over the long term. Here we use $\text{E}(R)$ to indicate the expectation of the random variable $R$. When the mean changes over time, such as in strongly seasonal regions, we explicitly indicate that by writing $\text{E}(R)$ as a function of time, $t$: $\text{E}(R|t) = \mu(t)$ (Fig. A1). If $\text{E}(R)$ changes over time and space, we write $\text{E}(R|t, u) = \mu(t, u)$. We indicate the estimated $\mu(t, u)$ by adding a caret on the symbol: $\hat{\mu}(t, u)$. Note, however, that the estimate of $\mu(t, u)$ and what values it can take depend on what distribution we assume $R$ follows. For a Gamma distribution, $\mu(t, u)$ can take any positive value, but if we use NDVI as a proxy for $R$, then $\mu(t, u)$ is necessarily within the interval $(-1, 1)$.

**Variance in resources, Var($R$)**

In viewing resources as a random variable, we not only obtain a formal framework for describing $\text{E}(R)$, but also the spread around $\text{E}(R)$, which we quantify with its variance (i.e.,

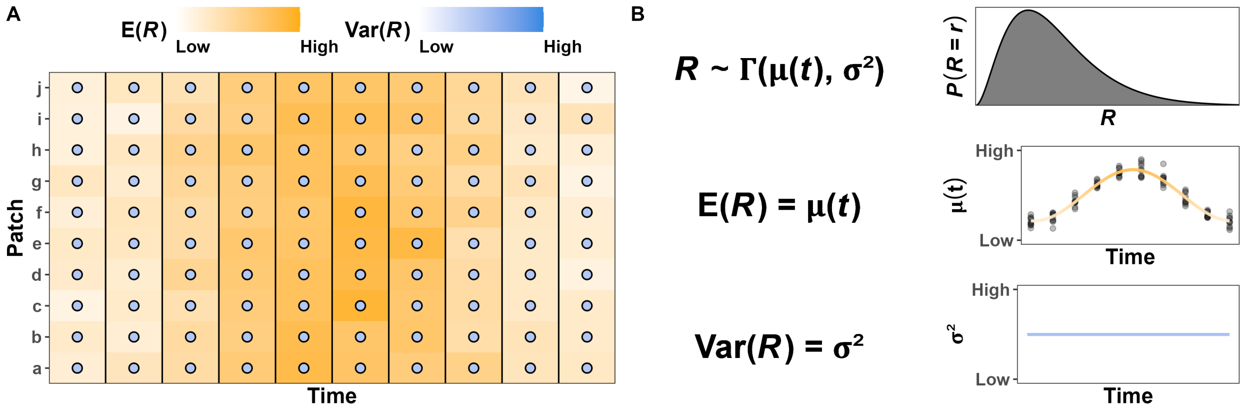


Figure A1: Fictitious example of variation in resources in a heterogeneous environment with constant variance around the mean. (A) Resources ($R$, orange fill) vary over time and space with a time-varying but spatially homogeneous mean ($\mu(t)$) and constant variance (dot color). (B) $R$ follows a Gamma distribution with mean $\mu(t)$ and constant variance $\mathrm{Var}(R) = \sigma^2$. The points in the central panel indicate the ralizations of $R$ in panel a.

stochasticity). We use the notation $\mathrm{Var}(R)$ to indicate the variance in $R$ around the mean, i.e., after accounting for changes in $\mu(t, u)$, and we use $\sigma^2(t, u)$ to indicate its function over time and time (with estimate $\widehat{\sigma^2}(t, u)$; see Fig. A2). For example, the time at which berry bushes produce fruit may seem highly stochastic (i.e., unpredictable) to a young bear, but it becomes predictable as it learns to understand and predict how the mean amount of berry changes with the seasons. Still, whether next year will be a good or bad year for berries remains stochastic because it depends on factors the bear cannot predict (e.g., droughts, fires), but the bear can still expect $\sigma^2(t, u)$ to be lower in the winter since berries will be hard to find (i.e., $\mu(t, u) \approx 0$). Also note that, like with $\mu(t, u)$, what values $\sigma^2(t, u)$ can take depends on what distribution we assume $R$ follows. Figures A3 and A4 show four examples of Gamma and Beta distributions, respectively, with different means and variances.

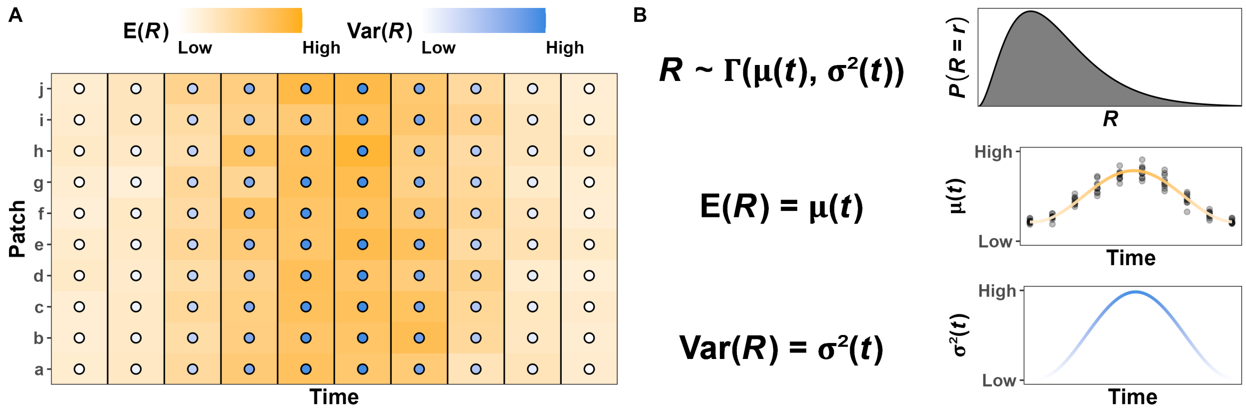


Figure A2: Fictitious example of variation in resources in a heterogeneous environment with changing variance around the mean. (A) Although resources ($R$, orange fill) vary over time and space, the variance (dot color) is lowest at the beginning and end of the observational period, and it is highest when $\mu(t)$ peaks. (B) $R$ follows a Gamma distribution with time-varying but spatially homogenous mean $\mu(t)$ and variance $\sigma^2(t)$. The points in the central panel indicate the ralizations of $R$ in panel a.

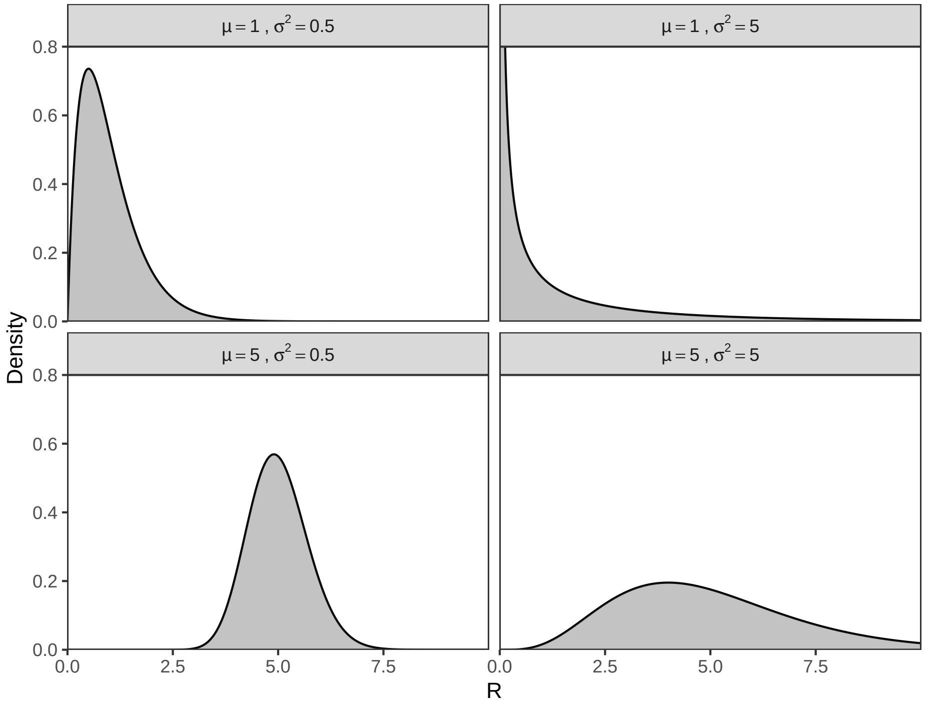


Figure A3: Examples of Gamma distributions with different means ($\mu$) and variances ($\sigma^2$).

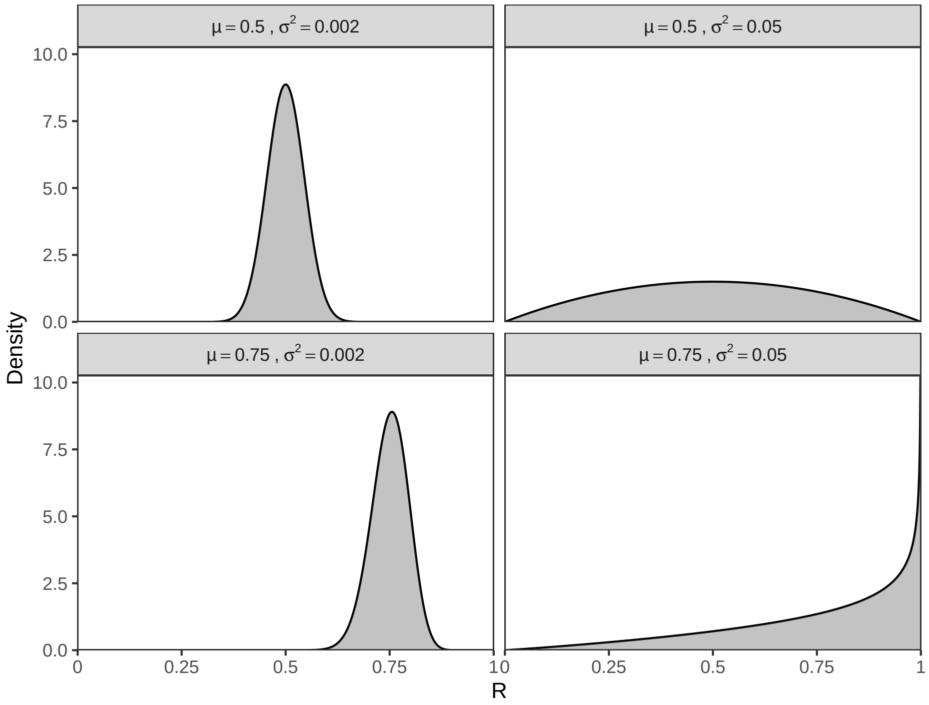


Figure A4: Examples of Beta distributions with different means ($\mu$) and variances ($\sigma^2$).
