## Appendix B: Simulations for "How resource abundance and stochasticity affect organisms’ range sizes"

### 1 Overview

This appendix illustrates all the steps necessary to produce the simulation used in the main manuscript (figs. 3 and 4). To achieve full transparency while minimizing computational times, the code illustrated in this pdf was not executed during the knitting of the document. Instead, the R Markdown document (`writing/appendix-2-simulations.Rmd`) contains code chunks that import the `RDS` files saved by the scripts used during the analysis via `R` code that is not printed in the pdf file. Although one can replicate the analyses by running the code in this pdf, we suggest only using this document for illustrative purposes. We suggest using the `R` scripts to replicate the simulations, instead.

### 2 Simulating the movement tracks

To reduce sampling variance between simulations at each time point in each panel of fig. 3 in the main text, we used the same set of 200 simulated tracks, which we generate in the `analysis/simulations/tracks.R` script. Most intermediate and diagnostic checks are not included in this document for the sake of brevity and simplicity, but their outputs and conclusions are listed in this section. In this appendix, we use the `ctmm` package (version 1.2.0, Fleming and Calabrese 2021) for movement modeling; the `terra` package (version 1.7-71, Hijmans 2024) to work with the rasters of simulated resources; the `dplyr` (version 1.1.4, Wickham et al. 2023), `purrr` (version 1.0.2, Wickham and Henry 2023), and `tidyr` (version 1.3.1, Wickham, Vaughan, and Girlich 2024) packages for data wrangling; and the the `ggplot2` (version 3.5.1, Wickham 2016), `cowplot` (version 1.1.3, Wilke 2024), and `gratia` (version 0.9.0, Simpson 2024) packages for plotting.

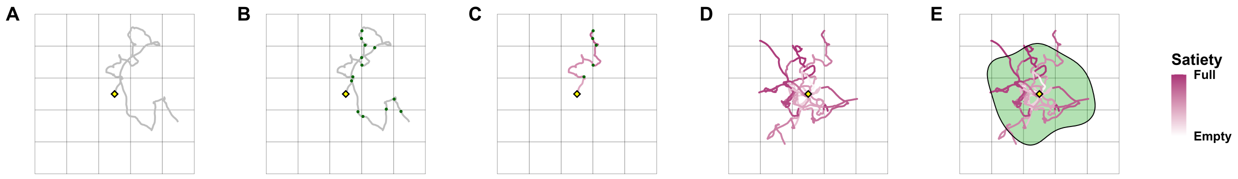


Figure B1: Overview of how organisms' home-range sizes were simulated. (a.) A track was simulated using an infinitely diffusive and continous-velocity movement model (Integrated Ornstein-Uhlenbeck, IOU), starting from the point $\langle 0, 0 \rangle$ (black and yellow square). (b.) Each time the track crossed into a new cell (green dots), the organism collected a random amount of resources that followed a Gamma distribution with mean $\mu(t)$ and variance $\sigma^2(t)$, with $t$ being fixed for the entirety of the track. (c.) Each time the organism collected more resources, its satiety (purple) increased. Once the organism collected sufficient resources, the organism stopped exploring (i.e., the track was truncated) and was allowed to return to $\langle 0, 0 \rangle$ over the same amount of time needed to reach satiety. (D) The process was repeated 200 times (13 tracks pictured in this panel) with the same value for $t$. (E) The final set of (truncated) tracks was then modeled using Ornstein-Uhlenbeck Foraging models to estimate the 95% home range size (see section 4).

```r
# NOTE: assuming the working directory is "hr-resource-stoch"
# attach all necessary packages
library('ctmm')    # for generating movement models and movement modeling
library('terra')   # for working with raster data
library('dplyr')   # for data wrangling
library('purrr')   # for functional programming
library('tidyr')   # for data wrangling (e.g., nested tibbles)
library('mgcv')    # for empirical Bayes GAMs
library('ggplot2') # for fancy plots
library('cowplot') # for fancy multi-panel plots
library('ggdag')   # for directed acyclical graphs
library('gratia')  # for ggplot-based GAM figures
theme_set(theme_bw()) # change default theme

# source custom functions
source('functions/rgamma2.R') # rgamma() parameterized by mean and variance
source('analysis/mean-variance-trends-panel-data.R') # mu and sigma2
source('analysis/simulations/movement-model.R') # for consistency
source('functions/get_hr.R') # for extracting gaussian home range
source('functions/label_visits.R') # decides when organism encounters food
source('analysis/figures/default-figure-styling.R') # for color palette

DELTA_T <- 60 # sampling interval in seconds
SAMPLES <- seq(0, 60 * 60 * 12, by = DELTA_T) # 12 hours by DELTA_T seconds

# projected raster of resources
PROJECTION <- '+proj=aeqd +lon_0=0 +lat_0=0 +datum=WGS84'
HABITAT <- matrix(data = 1, nrow = 500, ncol = 500) %>%
  raster(xmx = 1e3, xmn = -1e3, ymx = 1e3, ymn = -1e3, crs = PROJECTION)

# infinitely diffusive movement model
model <- ctmm(tau = c(Inf, 1e3), sigma = 0.1, mu = c(0, 0))

N_DAYS <- 2^10 # number of "days" (i.e., tracks with different seeds)

# extracts tracks from a ctmm movement model for given sample times
get_tracks <- function(day, times = SAMPLES) {
  simulate(model, # ctmm movement model
           t = times, # sampling times in seconds
           seed = day, # for a consistent track each day
           complete = TRUE, # add lat, long, and timestamp to telemetry
           crs = PROJECTION) # CRS projection string
}
```

```r
# generate simulated tracks (will be truncated at satiety later)
tels <- tibble(day = 1:N_DAYS, # a simulation for each day
               tel = map(.x = day, # set a seed for consistent results
                         .f = get_tracks)) # function to generate tracks
tels
```

```
# A tibble: 1,024 x 2
     day tel
   <int> <list>
 1     1 <telemtry[,8]>
 2     2 <telemtry[,8]>
 3     3 <telemtry[,8]>
 4     4 <telemtry[,8]>
 5     5 <telemtry[,8]>
 6     6 <telemtry[,8]>
 7     7 <telemtry[,8]>
 8     8 <telemtry[,8]>
 9     9 <telemtry[,8]>
10    10 <telemtry[,8]>
# i 1,014 more rows
```

```r
# find patch visits
tracks <- transmute(tels, # drop tel column
                    day, # keep day column
                    track = map(.x = tel, # add a column of full tracks
                                .f = \(x) {
                                  label_visits(.tel = x, .habitat = HABITAT)
                                }))
tracks <- tidyr::unnest(tracks, track) # make a single, large tibble
head(tracks)
```

```
# A tibble: 6 x 11
    day     t      x      y        vx      vy    longitude    latitude
  <int> <dbl>  <dbl>  <dbl>     <dbl>   <dbl>        <dbl>       <dbl>
1     1     0  0       0      0         0       0            0
2     1    60  0.0363 0.0593 -0.000529 0.00271  0.000000326 0.000000536
3     1   120  0.0722 0.395   0.000595 0.00755  0.000000648 0.00000358
4     1   180  0.0276 0.967  -0.00517  0.0102   0.000000248 0.00000875
5     1   240 -0.273  1.66   -0.00323  0.0130  -0.00000245  0.0000150
6     1   300 -0.351  2.42    0.000977 0.00894 -0.00000315  0.0000219
# i 3 more variables: timestamp <dttm>, cell_id <dbl>, new_cell <lgl>
```

After generating the tracks, we performed the following tests to ensure the number and length of the tracks were large enough for results to be stable. For the sake of conciseness, the code for each of the checks is not presented in this appendix, but it is available in the `R` scripts referenced in each section.

##### 2.1 Checking whether adding return trips is necessary

Script: `analysis/simulations/sensitivity-analyses/1a-return-sensitivity.R`

Adding return trips to $\langle 0, 0 \rangle$ after an organism reached satiety doubled computational times without appreciable improvements on the Gaussian home range estimates (including their 95% confidence intervals).

##### 2.2 Checking whether the sampling interval is sufficiently small

Script: `analysis/simulations/sensitivity-analyses/1b-delta-t-sensitivity.R`

Using three tracks generated with three arbitrary seeds (1, 2, and 3), we explored the effects of sampling interval ($\Delta t$) on the number of encounters with food (i.e., movements to new cells) detected. From each of the four checks, we created an exploratory plot that we present in figure B2.

**Exploratory plot B2A.** The amount of time between encounters ranged from 1 second (the minimum $\Delta t$) to 24 minutes and 10 seconds. Approximately 93% of the encounters (500/536, excluding the first 3 events of each track) occurred with 30 or more seconds between events.

**Exploratory plot B2B.** Halving the sampling interval had little to no effect on the total number of encounters for $\Delta t \lessapprox 60\,s$.

**Exploratory plot B2C.** A sampling interval of $\Delta t = 30$ seconds was small enough to capture fine-scale movement in the tracks but large enough to avoid excessive amounts of data and an inflated amount of encounters when an organism was near cell boundaries.

**Exploratory plot B2D.** The three tracks used for these exploratory plots are sufficiently different that we considered them to be a representative sample of movement tracks simulated by the OUF model. All three tracks have a reasonable amount of both tortuous and directed movement.

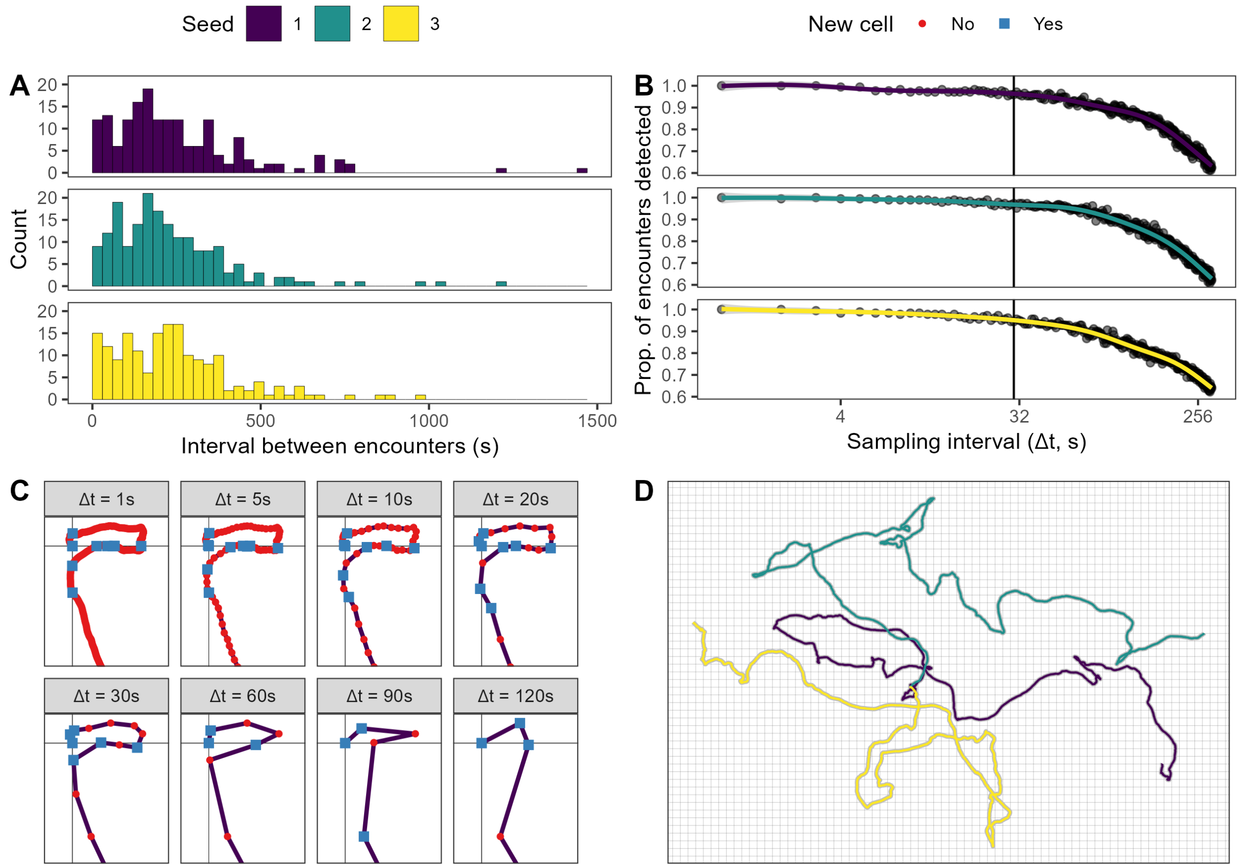


Figure B2: Exploratory plots used to decide an appropriate sampling interval. (A) Histograms of the number of encounters as a function of the interval between encounters, with a binwith of 30 seconds. Although some encounters occur with less than 30 seconds between them, approximately 93% of them occur at least 30 seconds apart. (B) Number of detected encounters as a function of sampling interval. The colored lines indicate the estimated relationship based on a Generalized Additive Model fit using the `geom_smooth` function from the `ggplot2` package. Although the number of encounters detected decreases as sampling interval doubles, the loss at $\Delta = 30$s is negligible (note that all sampling times included $>$ 60% of the encounters). (C) Beginning of the track generated with seed "1" (purple line) for different sampling intervals. Red dots indicate locations where the organism remained in the same cell, while the blue squares indicate when an organism was in a new cell and thus encountered food. While the number of encounters detected decreases as the sampling interval increases, most of the encounters lost at $\Delta t = 30$s occured because the organism remained almost adjacent to the borders between cells. Additionally, the track at $\Delta t = 30$s is still sufficiently tortuous to represent realistic movement. (D) The three tracks used in these tests over the raster used for determining when the organism encountered food.

##### 2.3 Checking how many tracks were necessary

As the organism moved, it encountered $R$ resources sampled from a family of Gamma distributions with mean $\mu(t)$ and variance $\sigma^2(t)$. The figure below shows the range in Gamma distributions for the minimum, median, and maximum $\mu(t)$ and $\sigma^2(t)$.

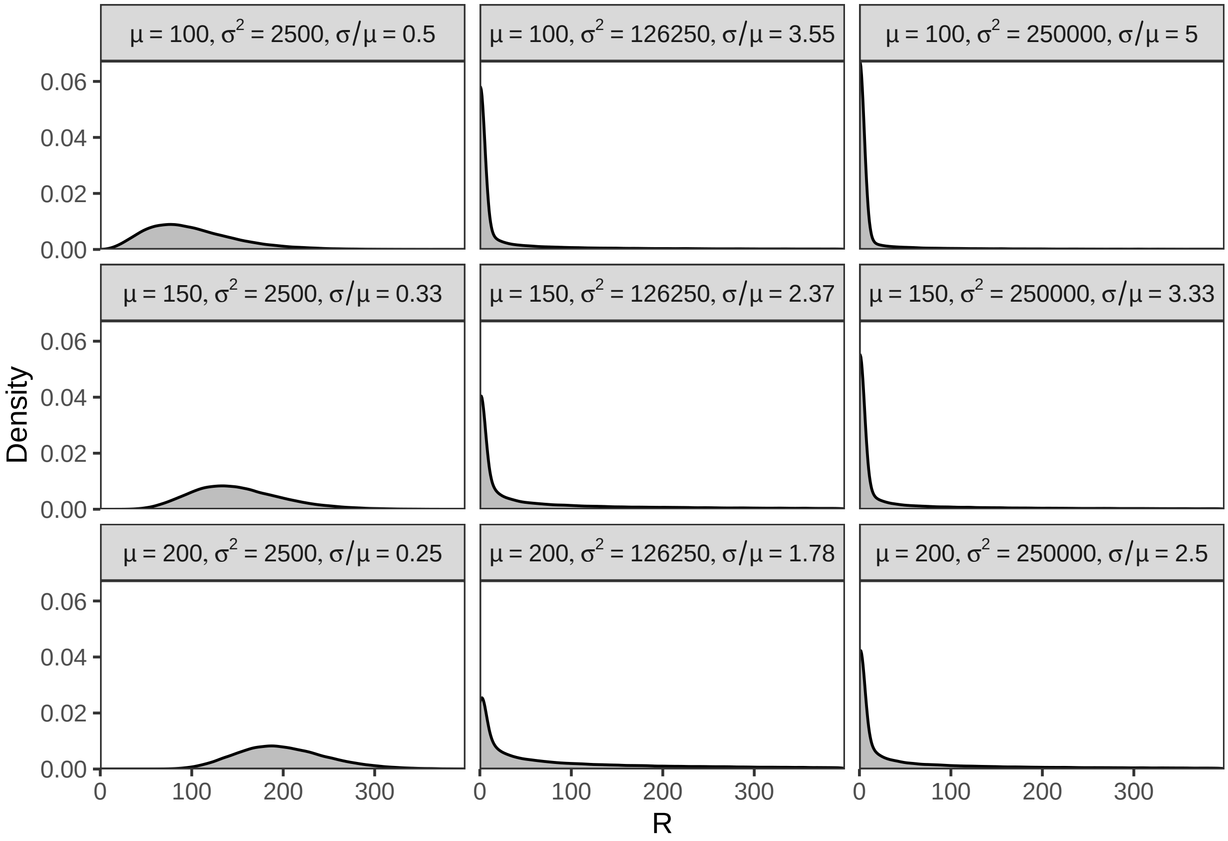


Figure B3: Gamma distributions for the minimum, median, and maximum mean (100, 150, 200) and variance (2 500, 126 250, 250 000) with the corresponding coefficients of variation. The medians and modes of the distributions increase with the mean but decrease with the variance. Greater coefficients of variation result in wider distributions with greater density near zero.

Script: `analysis/simulations/2-hr-mean-variance-simulations-days.R`

The end of this script generates figures that show the mean time spent searching for resources before reaching satiety. This appendix includes the two figures that show the change in time required to reach satiety as a function of the number of tracks. The estimated times for each number of replicates can be found in the `figures/5-by-5-sensitivity-analysis` folder.

```r
all <-
  tibble(d = map_dfr(
    list.files('simulations', pattern = '*-replicates.rds',
               full.names = TRUE),
    function(fn) {
      readRDS(fn) %>%
        mutate(mean = paste(mean, 'mean'),
               variance = paste(variance, 'var'),
               replicates = max(day)) %>%
        group_by(mean, variance, animal, replicates) %>%
        summarize(est = mean(t_expl),
                  .groups = 'drop')
    })) %>%
  unnest(d) %>%
  mutate(est = est / (1 %#% 'hours')) %>%
  pivot_wider(names_from = replicates, values_from = est) %>%
  pivot_longer(cols = as.character(2^(4:9)), values_to = 'est',
               names_to = 'replicates') %>%
  mutate(diff_est = est - `1024`,
         rel_est = est / `1024`,
         replicates = factor(replicates, levels = 2^(4:9)))

ggplot(all) +
  facet_wrap(~ replicates) +
  geom_line(aes(animal, diff_est, group = paste(mean, variance)),
            alpha = 0.1) +
  geom_hline(yintercept = 0, color = 'red') +
  labs(x = 'Time', y = 'Difference in average daily time required (hours)')
```

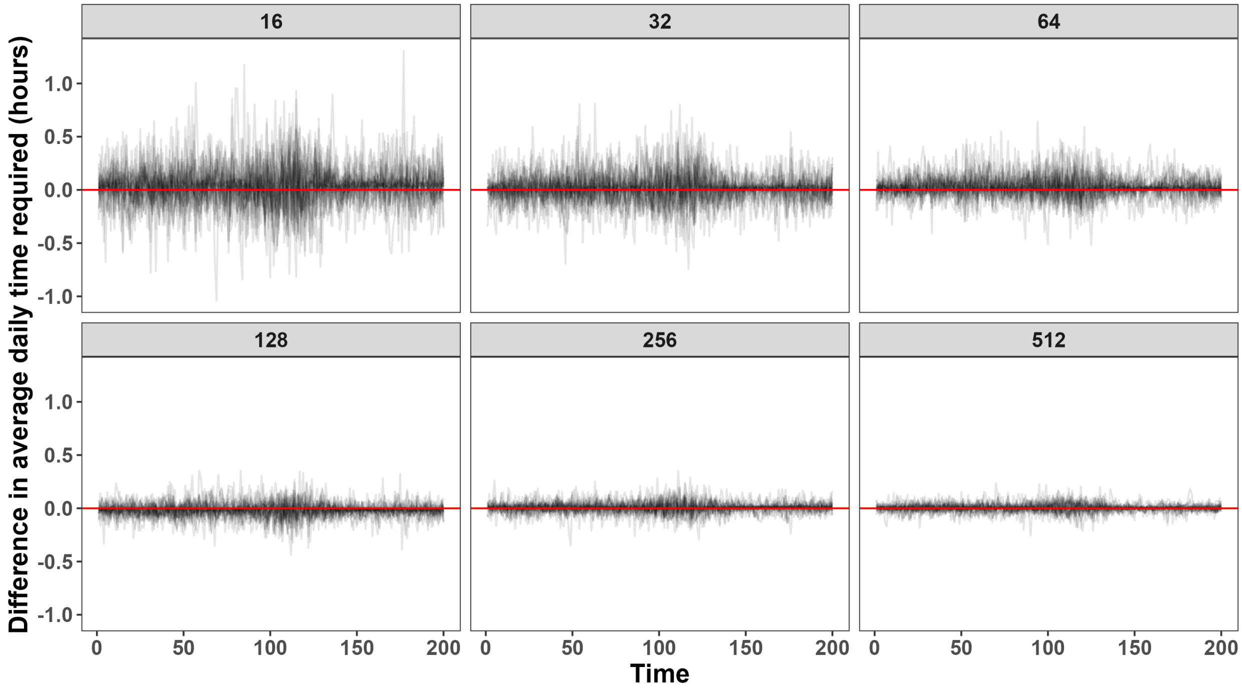


Figure B4: Difference in average time required to reach satiety for each scenario in figure 1 (indicated by each line) and different amounts of tracks (indicated at the top of each facet). The differences are relative to simulations with $2^{10} = 1024$ tracks for each time point on the x axis (and each scenario in figure 1). The amount of time stabilizes for $\gtrapprox$ 100 tracks, which suggests that the home range estimates should also be stable with more than 100 tracks.

```r
ggplot(all) +
  facet_wrap(~ replicates) +
  geom_line(aes(animal, rel_est, group = paste(mean, variance)),
            alpha = 0.1) +
  geom_hline(yintercept = 1, color = 'darkorange') +
  scale_y_continuous(
    trans = 'log2', breaks = 2^seq(-1, 1, by = 0.5),
    labels = parse(text = paste0('2^', seq(-1, 1, by = 0.5))))+
  labs(x = 'Time',
       y = expression(bold(paste('Relative difference in time to satiety'~
                                 (log[2]~scale)))))
```

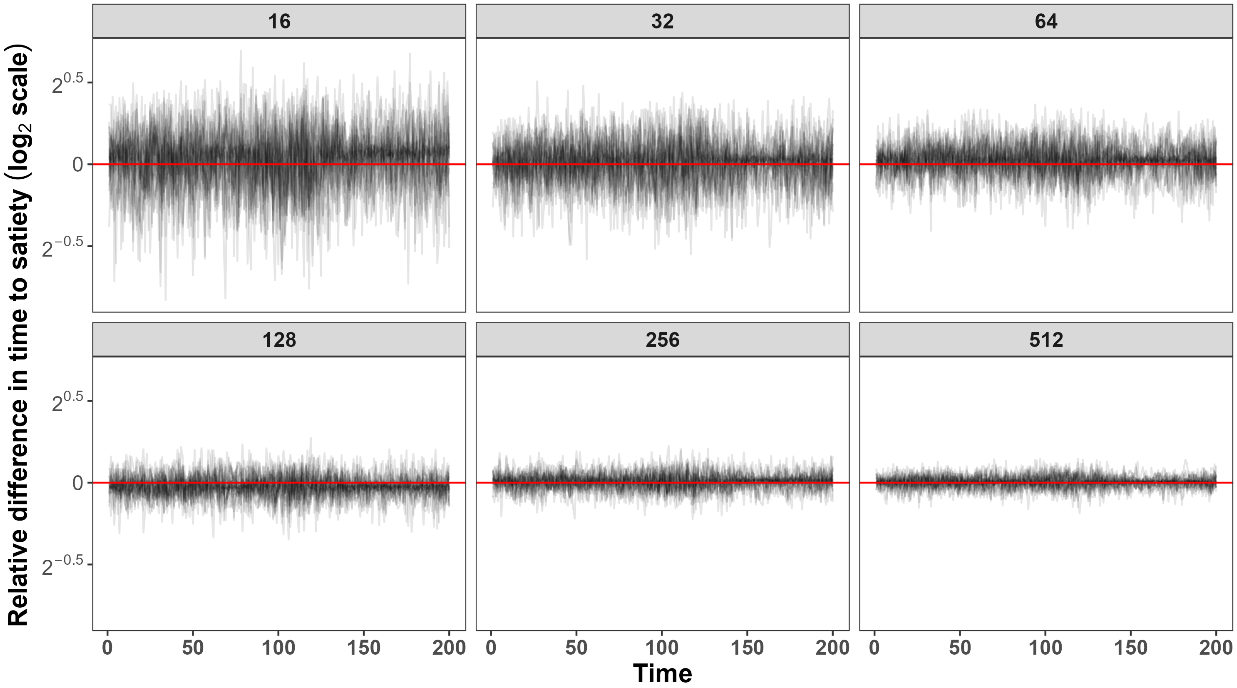


Figure B5: Ratios of average time required to reach satiety for each scenario in figure 1 (indicated by each line) and different amounts of tracks (indicated at the top of each facet). The ratios are relative to simulations with $2^{10} = 1024$ tracks for each time point on the x axis (and each scenario in fig. 1). The amount of time stabilizes for $\gtrapprox 100$ tracks, which suggests that the home range estimates should also be stable with more than 100 tracks.

Script: `analysis/simulations/sensitivity-analyses/1c-hr-simulation-extreme-scenarios.R`

In this script, we check how many tracks are necessary to produce stable and accurate home-range size estimates. We do so by estimating the home-range sizes from varying numbers of track in the best-case scenario (highest $\mathrm{E}(R)$ and lowest $\mathrm{Var}(R)$) and worst-case scenario (lowest $\mathrm{E}(R)$ and highest $\mathrm{Var}(R)$).

```r
set.seed(1) # for consistent results
tels <- readRDS('simulations/tracks.rds') # list of telemetry tracks
tracks <- readRDS('simulations/labelled-tracks.rds') # tibble of tracks
MAX_T <- max(tracks$t) # maximum amount of exploration time

WORST <- filter(d55, mu == min(mu)) %>% # lowest mean resources
  filter(sigma2 == max(sigma2)) %>% # with highest variance
  slice(1) # take the first row only
BEST <- filter(d55, mu == max(mu)) %>% # highest mean resources
  filter(sigma2 == min(sigma2)) %>% # with lowest variance
  slice(1) # take the first row only
```

```r
days <-
  transmute(bind_rows(WORST, BEST),
            animal,
            mu,
            sigma2,
            d = list(tracks),
            scenario = c('Worst case', 'Best case')) %>%
  unnest(d) %>% # unnest the datasets so we have a single, large tibble
  select(-timestamp) %>%
  # generate the food for each row from a gamma distribution
  mutate(food = rgamma2(mu = mu, sigma2 = sigma2, N = n()),
         # the animal finds food if it visits a new cell, otherwise not
         food = if_else(new_cell, food, 0)) %>%
  # end the movement once the animal has reached satiety
  group_by(day, animal, scenario) %>%
  # calculate the total visits, total calories, and if animal is full
  mutate(satiety = cumsum(food), # for diagnostics if animals aren't full
         full = satiety >= REQUIRED) %>% # did the animal reach its needs?
  filter(cumsum(full) <= 1) %>% # full only once
  ungroup()

# single estimates that eventually converge to the asymptote ----
days_summarized <-
  days %>%
  # find how long it took to reach satiety
  group_by(scenario, day) %>%
  nest(tel_day = -c(scenario, day)) %>%
  mutate(t_expl = map_dbl(tel_day, \(d) max(d$t))) %>%
  # add days sequentially
  group_by(scenario) %>%
  mutate(t_start = lag(2 * t_expl), # add the return time before next "day"
         t_start = if_else(is.na(t_start), 0, t_start), # start at 0
         t_start = cumsum(t_start), # make start times comsecutive
         tel_day = map2(day, t_expl,
                        \(i, te) tels$tel[[i]] %>% # extract day's tel
                          data.frame() %>% # for filtering
                          filter(t <= te))) %>% # end tracks at satiety
  unnest(tel_day) %>% # make one big dataset
  mutate(t = t + t_start, # make times consecutive
         individual.local.identifier = scenario, # ctmm identifier
         timestamp = as.POSIXct(t, origin = '2000-01-01')) %>% # new times
  ungroup() # remove grouping by scenario
```

```r
# estimate saturation curve of home range size over number of days
saturation_days <-
  expand_grid(n_days = (2^seq(1, log2(1e3), by = 0.2)) %>%
                round() %>%
                unique(),
              case = unique(days_summarized$scenario)) %>%
  mutate(data = map2(n_days, case,
                     \(.n, .case) filter(days_summarized,
                                         day <= .n,
                                         scenario == .case)),
         tel = map(data, as.telemetry), # convert to telemetry for modeling
         theta = map(tel, \(x) ctmm.guess(data = x, interactive = FALSE)),
         m = map(1:n(), \(i) {
           cat('Fitting model', i, '\n')
           ctmm.fit(tel[[i]], theta[[i]])
         }), # fit movement model
         sigma = map_dbl(m, \(.m) ctmm:::area.covm(.m$sigma)), # var(pos)
         hr = get_hr(.sigma = sigma, quantile = 0.95)) # Gaussian HR

saturation_days %>%
  select(case, n_days, sigma, hr) %>%
  readr::write_csv('simulations/hr-saturation-days.csv')

ggplot(saturation_days, aes(n_days, hr)) +
  facet_wrap(~ case, nrow = 1) +
  geom_vline(xintercept = 200, color = 'darkorange') +
  geom_smooth(method = 'gam', color = 'black',
              formula = y ~ s(x, bs = 'cs', k = 10),
              method.args = list(family = Gamma(link = 'log'))) +
  geom_point(alpha = 0.3) +
  scale_x_continuous(expression(Number~of~days~sampled~(log[2]~scale)),
                     trans = 'log2', breaks = c(2, 16, 128, 1024),
                     limits = c(2, 1100)) +
  scale_y_continuous(expression(atop(Estimated~'home-range',
                                     size~(log[2]~scale))),
                     trans = 'log2')
```

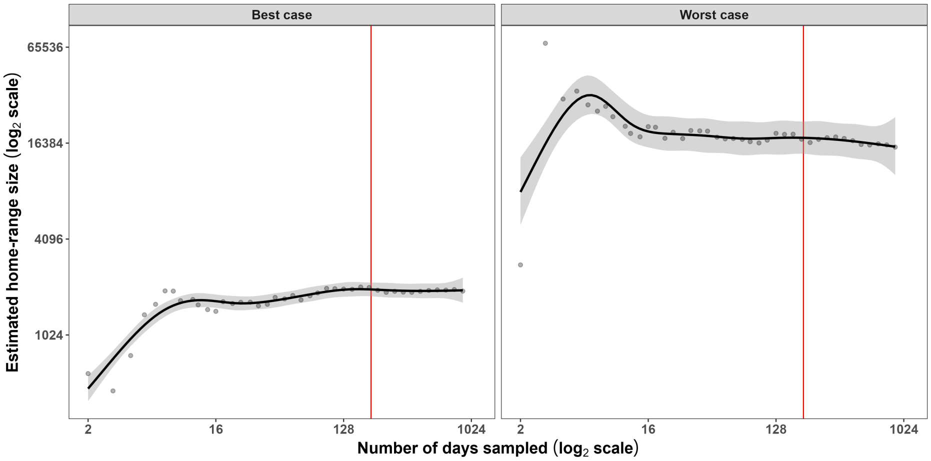


Figure B6: Estimated home-range sizes as a function of the number of days sampled for an organism in a habitat with the highest E($R$) and lowest Var($R$) (left) and an organism with the lowest E($R$) and highest Var($R$) (right). In both cases, 200 days are sufficient to produce stable estimates of home-range size.

### 3 Main scripts (to be run in the following order)

1. `analysis/simulations/hr-mean-variance-simulations-days.R`
2. `analysis/simulations/hr-mean-variance-simulations-days-summarized.R`
3. `analysis/simulations/hr-mean-variance-simulations-modeling.R`
4. `analysis/simulations/hr-mean-variance-simulations-hrs.R`
5. `analysis/simulations/modeling-R-and-hr.R`

### 4 Modeling the results from the simulations

This final section illustrates the location-scale GAM used to estimate the effects of resource abundance and stochasticity on the simulated home-range sizes. Although the model was fit to both the 95% and core (50%) home range estimates, the results were presented only for the 95% quantile for simplicity. The home range was caulculated using a Gaussian approximation and the simulated organism's positional variance via the formula:

$$\widehat{h_q} = -2\log(1-q)\pi\varsigma^2,$$

where $\widehat{h_q}$ is the estimated home range for the $q^{\text{th}}$ quantile and $\varsigma^2$ is the positional variance. Consequently, the estimated core home range estimate is simply a multiple of the 95% quantile:

$$\widehat{h_{0.95}}/\widehat{h_{0.5}} = \frac{-2\log(1-0.95)\pi\sigma^2_{pos}}{-2\log(1-0.5)\pi\sigma^2_{pos}} = \frac{\log(1-0.95)}{\log(1-0.5)} = \frac{\log(0.05)}{\log(0.5)} \approx 4.322,$$

and therefore $\mathrm{E}(R)$ and $\mathrm{Var}(R)$ would have similar effects on the core home range as on the 95% home range (but about 4.322 times weaker). Note that we used this approximation because, unlike real organisms, the simulated organism did not use the available space selectively (e.g., spending more time near cell boundaries), so the Gaussian approximation is appropriate. When estimating home-range sizes based on empirical data, we suggest estimating utilization distributions using methods such as Autocorrelated Kernel Density Estimation (see Appendix C and: Noonan et al. 2019; Alston et al. 2022; Silva et al. 2022).

```r
# switch data to long format with a column of quantile
sims <- readRDS('H:/GitHub/hr-resource-stoch/simulations/days-hrs.rds') %>%
  tibble() %>%
  pivot_longer(c(hr_50, hr_95), names_to = 'quantile', values_to = 'hr') %>%
  mutate(quantile = factor(quantile)) # necessary for GAMs
sims
```

```
# A tibble: 10,000 x 9
   mean     variance animal    mu sigma2   t_expl pos_var quantile    hr
   <fct>    <fct>     <int> <dbl>  <dbl>    <dbl>   <dbl> <fct>    <dbl>
 1 constant constant      1   150 126250  8738220    287. hr_50    1250.
 2 constant constant      1   150 126250  8738220    287. hr_95    5404.
 3 constant constant     10   150 126250  8803140    282. hr_50    1229.
 4 constant constant     10   150 126250  8803140    282. hr_95    5312.
 5 constant constant    100   150 126250  8710920    281. hr_50    1224.
 6 constant constant    100   150 126250  8710920    281. hr_95    5288.
 7 constant erratic     200   150   2500  7366620    162. hr_50     706.
 8 constant erratic     200   150   2500  7366620    162. hr_95    3051.
 9 linear   constant      1   100 126250 14073060    631. hr_50    2747.
10 linear   constant      1   100 126250 14073060    631. hr_95   11873.
# i 9,990 more rows
```

The sampling of the data is not uniform, but it is not particularly problematic.

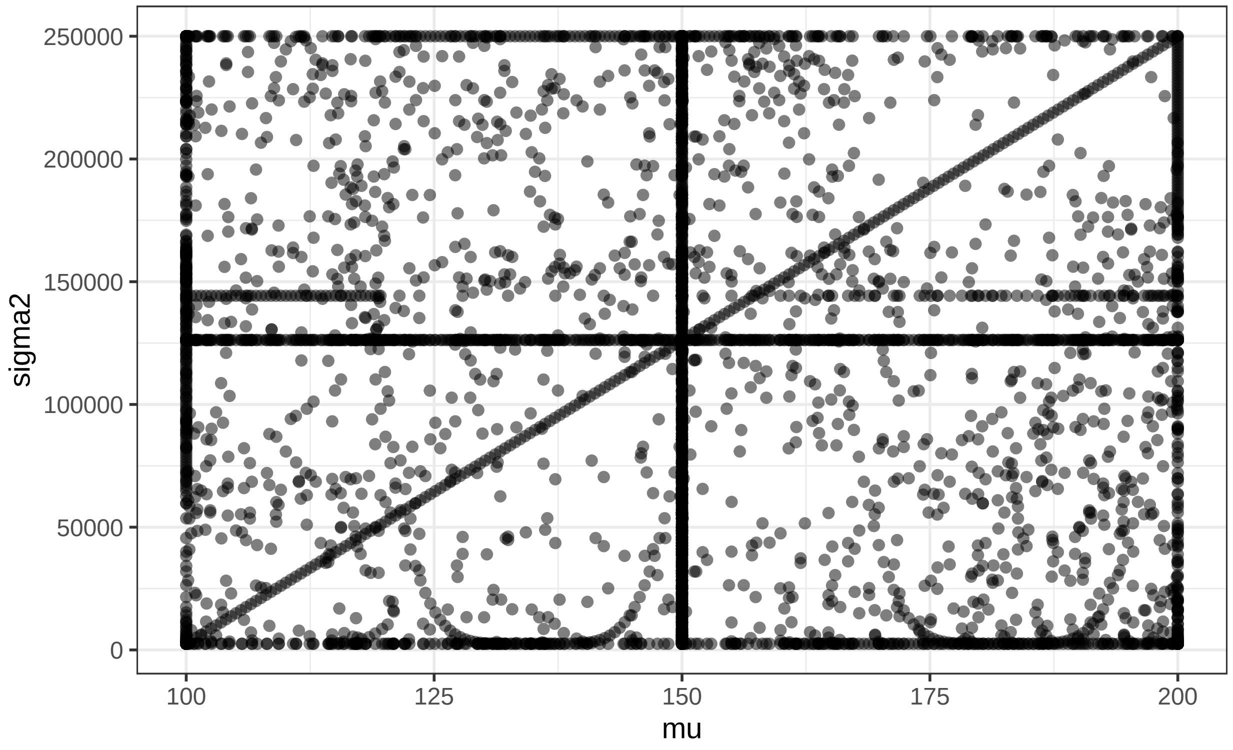


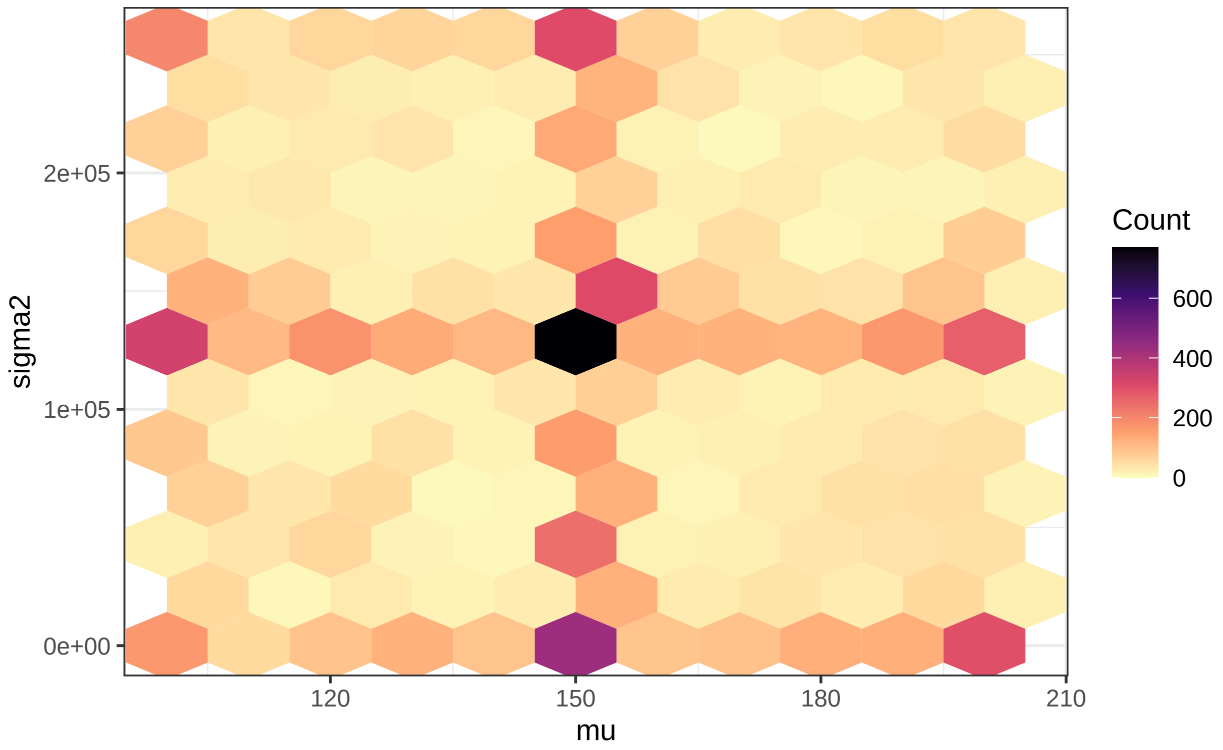

The abundance of data for the average $\mu(t)$ and $\sigma^2(t)$ (see figures above) did not create biases in the models, since removing such values had no appreciable effect on the model estimates (not shown). We estimated the effects of $\mathrm{E}(R)$ and $\mathrm{Var}(R)$ on the simulated home-range sizes using a location-scale GAMs (GAMLS) with a gamma family of distributions (since home-range sizes are strictly positive). We hypothesized that the effect of $\mathrm{Var}(R)$ would depend on $\mathrm{E}(R)$, since we expected organisms to respond to changes in $\sigma^2(t)$ less when resources were abundant than when they were scarce (fig. B7).

Within a Bayesian framework, Directed Acyclical Graphs (DAGs) can be used to infer causality (McElreath 2016). The DAG below can be interpreted as follows: $\mathrm{E}(R)$ and $\mathrm{Var}(R)$ are the (controlled) variables that determine resources, $R$, and, consequently, an organism's home-range size, $H$. Additionally, as indicated by the arrow going from $\mathrm{E}(R)$ to $\mathrm{Var}(R)$, the effect of $\mathrm{Var}(R)$ depends of the value of $\mathrm{E}(R)$ (and vice-versa, but arrows in DAGs have to be unidirectional).

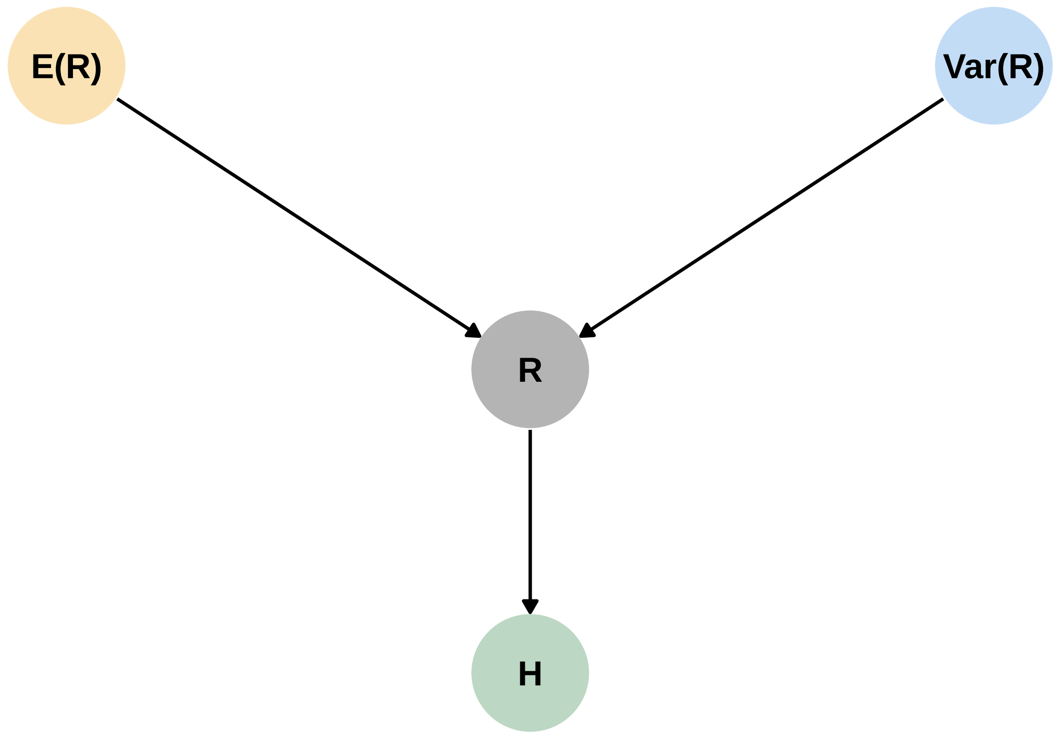


Figure B7: Directed Acyclical Graph assumed for inferring the causal effects of $\mathrm{E}(R)$ and $\mathrm{Var}(R)$ on $H$.

```
m <- gam(
  list(
    hr ~ quantile + s(mu, k = 5) + s(sigma2, k = 5) + ti(mu, sigma2, k = 5),
    ~ quantile + s(mu, k = 5) + s(sigma2, k = 5) + ti(mu, sigma2, k = 5)),
  family = gammals(),
  data = sims,
  method = 'REML')
```

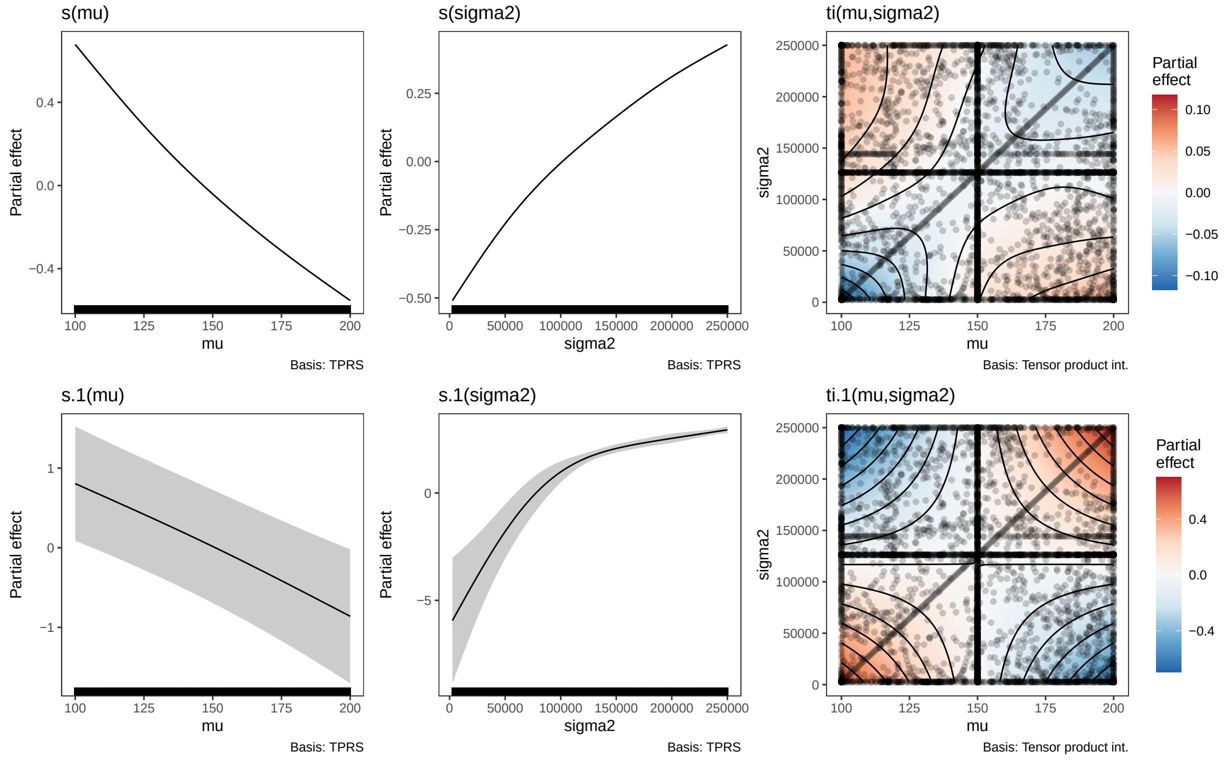


Figure B8: Marginal and interaction effects of each of the terms present in the final GAMLS (on the log link scale). The shaded ribbons indicate the 95% Bayesian credible intervals. The data are included as rug plots in the marginal terms and as points in the interaction terms.

```r
summary(m)
```

```

Family: gammals
Link function: identity log

Formula:
hr ~ quantile + s(mu, k = 5) + s(sigma2, k = 5) + ti(mu, sigma2,
    k = 5)
~quantile + s(mu, k = 5) + s(sigma2, k = 5) + ti(mu, sigma2,
    k = 5)

Parametric coefficients:
                  Estimate Std. Error   z value Pr(>|z|)
(Intercept)      7.097e+00  4.949e-04 14340.603  < 2e-16 ***
quantilehr_95    1.464e+00  6.848e-04  2137.506  < 2e-16 ***
(Intercept).1   -2.875e+00  3.636e-01    -7.907 2.63e-15 ***
quantilehr_95.1  1.492e-13  9.673e-02     0.000        1
---
Signif. codes:  0 '***' 0.001 '**' 0.01 '*' 0.05 '.' 0.1 ' ' 1

Approximate significance of smooth terms:
                   edf Ref.df    Chi.sq  p-value
s(mu)            3.990  4.000 1.074e+06  < 2e-16 ***
s(sigma2)        3.995  4.000 7.752e+05  < 2e-16 ***
ti(mu,sigma2)   14.325 15.662 8.625e+03  < 2e-16 ***
s.1(mu)          1.309  1.542 2.753e+01 3.09e-06 ***
s.1(sigma2)      3.404  3.739 1.166e+02  < 2e-16 ***
ti.1(mu,sigma2)  1.032  1.060 9.571e+00  0.00219 **
---
Signif. codes:  0 '***' 0.001 '**' 0.01 '*' 0.05 '.' 0.1 ' ' 1

Deviance explained = 99.9%
-REML =  57981  Scale est. = 1         n = 10000
```

Fig. B8 shows each of the terms from the model. The estimated effect of `quantile` (see the output of `summary()` above) supports the relationship between the gaussian HR estimates detailed above, since $\exp(1.464) = 4.323 \approx \frac{\log 0.5}{\log 0.05}$, but there is no differences in the scale parameter between the two quantiles.

McElreath, Richard. 2016. *Statistical Rethinking: A Bayesian Course with Examples in r and Stan*. Chapman & Hall/CRC Texts in Statistical Science Series 122. Boca Raton: CRC Press/Taylor & Francis Group.

Noonan, Michael J., Marlee A. Tucker, Christen H. Fleming, Thomas S. Akre, Susan C. Alberts, Abdullahi H. Ali, Jeanne Altmann, et al. 2019. "A Comprehensive Analysis of Autocorrelation and Bias in Home Range Estimation." *Ecological Monographs* 89 (2): e01344. https://doi.org/10.1002/ecm.1344.

Silva, Inês, Christen H. Fleming, Michael J. Noonan, Jesse Alston, Cody Folta, William F. Fagan, and Justin M. Calabrese. 2022. "Autocorrelation-informed Home Range Estimation: A Review and Practical Guide." *Methods in Ecology and Evolution* 13 (3): 534–44. https://doi.org/10.1111/2041-210X.13786.

Simpson, Gavin L. 2024. *Gratia: Graceful Ggplot-Based Graphics and Other Functions for GAMs Fitted Using Mgcv*. https://gavinsimpson.github.io/gratia/.

Wickham, Hadley. 2016. *Ggplot2: Elegant Graphics for Data Analysis*. Springer-Verlag New York. https://ggplot2.tidyverse.org.

Wickham, Hadley, Romain Francois, Lionel Henry, Kirill Muller, and Davis Vaughan. 2023. *Dplyr: A Grammar of Data Manipulation.* https://CRAN.R-project.org/package=dplyr.

Wickham, Hadley, and Lionel Henry. 2023. *Purrr: Functional Programming Tools.* https://CRAN.R-project.org/package=purrr.

Wickham, Hadley, Davis Vaughan, and Maximilian Girlich. 2024. *Tidyr: Tidy Messy Data.* https://CRAN.R-project.org/package=tidyr.

Wilke, Claus O. 2024. *Cowplot: Streamlined Plot Theme and Plot Annotations for 'Ggplot2'.* https://CRAN.R-project.org/package=cowplot.
