## Appendix C: Empirical modeling for "How resource abundance and stochasticity affect organisms’ range sizes"

#### Contents


**1 Modeling $R$** **2**

**2 Estimating $R$ using NDVI** **3**

**3 Reproducing the analyses** **4**

**4 Modeling the tapir's movement over time** **6**

**5 Modeling $\mathbf{E}(R)$ and $\mathbf{Var}(R)$ over time** **8**

**6 Modeling the effects of $\mathbf{E}(R)$ and $\mathbf{Var}(R)$ on space use** **13**

**References** **17**

### 1 Modeling $R$

Location-scale models (theory: Rigby and Stasinopoulos 2005; Stasinopoulos and Rigby 2007; examples: Bjorndahl et al. 2022; Mariën et al. 2022; Gushulak et al. 2024) are a class of statistical models that allow us to estimate changes in a random variable's mean (i.e. its location) and variance (which depends on its scale) while allowing the mean-variance relationship to vary. `mgcv` (Wood 2017) is a commonly used package for `R` (R Core Team 2023) that allows one to fit location-scale models with various families of distributions, including Gaussian (i.e., normal), gamma, and Tweedie location-scale families. The Gaussian location-scale family is very flexible, since the mean and variance parameters are assumed to be independent, but it is inappropriate for strictly positive data (e.g. available biomass) and other bounded data (e.g., proportions and Normalized Difference Vegetation Index, i.e. NDVI, see Pettorelli et al. 2005; Pettorelli et al. 2011). The Gamma location-scale family is best for strictly positive responses, such as elemental compositions (e.g., carbon to nitrogen ratio, see Rizzuto et al. 2021), total biomass, or energetic intake. The Tweedie location-scale family is similar to the Gamma family, but it allows for zero data, so it is appropriate for data with a non-trivial amount of zeros, such as daily precipitation or prey density (but see zero-inflated distributions: Zuur et al. 2009). For this paper, we estimated $R$ by modeling NDVI using `mgcv` and a beta location-scale family. While the family is not available in `mgcv` at the time of publication, the code for it is avialable on GitHub at https://github.com/QuantitativeEcologyLab/hr-resource-stoch/blob/main/functions/betals.r. If one is interested in families of distributions which are not available in `mgcv`, we suggest using the `brms` package (Bürkner 2017), which supports fully distributional, Bayesian models (Bürkner 2018).

Modeling the mean and variance terms of $R$ should be done carefully. Since trends in both $\mathrm{E}(R)$ and $\mathrm{Var}(R)$ can be spatiotemporally nonlinear and non-monotonic, we suggest using a GAM rather than a GLM. However, the complexity of the spatiotemporal terms should be chosen carefully, particularly for the mean's terms. An excessively wiggly $\hat{\mu}(t,u)$ will cause $\sigma^2(t,u)$ to be under-estimated, while an excessively smooth $\hat{\mu}(t,u)$ will cause $\sigma^2(t,u)$ to be over-estimated. Although there is no error-proof system, choosing the complexity of the terms based on the organism's ability to detect change and adapt is a reasonable starting point. Additionally, using restricted marginal likelihood (`method = 'REML'`, see Wood 2011) should help constrain the complexity of the smooths. Simpson (2018) provides a useful introduction to GAMs for biological time series.

#### 2 Estimating $R$ using NDVI

Since all NDVI values in our dataset were sufficiently greater than 0 (fig. C1), we defined $R$ as following a spatiotemporally-varying beta distribution with mean $\mu(t,u)$ and variance $\sigma^2(t,u)$: $R \sim B(\mu(t,u), \sigma^2(t,u))$. We use this parameterization here for ease of explanation, but note that beta distributions are generally parameterized using the shape parameters $\alpha$ and $\beta$ such that the mean is

$$\mathrm{E}(R) = \frac{\alpha}{\alpha + \beta} \tag{1}$$

while the variance is

$$\mathrm{Var}(R) = \frac{\alpha\beta}{(\alpha+\beta)^2(\alpha+\beta+1)}. \tag{2}$$

If NDVI values are near or below zero (e.g., in barren or snowy ecosystems), we suggest using the equation

$$\nu^* = \frac{\nu + 1}{2}, \tag{3}$$

where $\nu$ is the NDVI values in the [-1, 1] scale and $\nu^*$ is the NDVI values scaled to [0, 1]. Since the transformation is linear (i.e., it only involves addition and division), estimates of $\mathrm{E}(\nu^*)$ and $\mathrm{Var}(\nu^*)$ can be back-transformed to the [-1, 1] scale with no bias, unlike with nonlinear transformations such as $\arcsin\sqrt{\nu}$ and $\log(\nu + 1)$ (Jensen 1906; Denny 2017).

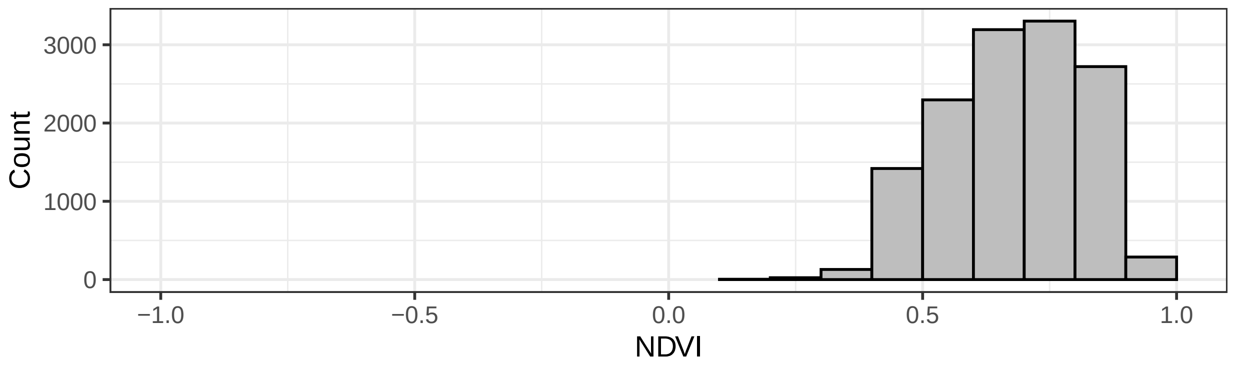


Figure C1: Histogram of the NDVI values used for the beta location-scale model (after removing the problematic raster for 2017-12-19; see section 5). Note that all values are far from zero (range: 0.3534 to 0.9475).

### 3 Reproducing the analyses

This section illustrates the steps necessary to reproduce the tapir movement analysis and the related figure in the manuscript (fig. 5). The tapir data used here is from the work of Medici et al. (2022) and can be found at the GitHub repository located at https://github.com/StefanoMezzini/tapirs. To minimize the computational costs of creating this appendix, we load the necessary objects through hidden `R` chunks rather than re-running all the code. Still, those interested in replicating the analyses can do so by using the code in the pdf document or the related `R` Markdown (`Rmd`) document (as well as the `R` scripts). All the packages and source scripts required to run the analyses in this document are listed in the code chunk below. For spatial data, we use the `MODIStsp` package (version 2.1.0, Busetto and Ranghetti 2016) to download the NDVI rasters, the `terra` package (version 1.7-71, Hijmans 2024) to work with the NDVI rasters, and the `sf` package (version 1.0-16, Pebesma 2018; Pebesma and Bivand 2023) to work with simple features (e.g., telemetry data and shapefiles). We use the `dplyr` (version 1.1.4, Wickham et al. 2023), `purrr` (version 1.0.2, Wickham and Henry 2023), and `tidyr` (version 1.3.1, Wickham et al. 2024) packages for data wrangling, and the `lubridate` package (version 1.9.3, Grolemund and Wickham 2011) for converting calendar dates to decimal dates. Finally, we used the `ctmm` package (version 1.1.0, Fleming and Calabrese 2021) and the `mgcv` package (version 1.9-1, Wood 2017) for modeling, and the `ggplot2` (version 3.5.1, Wickham 2016) and `cowplot` (version 1.1.3, Wilke 2024) packages for plotting. We start by attaching all the packages and custom functions we need for the following sections.

```
# NOTE: assuming the working directory is "hr-resource-stoch/writing"
library('terra')     # to import and save rasters
library('dplyr')     # for data wrangling
library('purrr')     # for functional programming
library('tidyr')     # for data wrangling
library('ggplot2')   # for fancy plots
library('cowplot')   # for fancy multi-panel plots
library('ctmm')      # for movement modeling
library('mgcv')      # for empirical Bayesian GAMs
library('lubridate') # for smoother date wrangling
library('sf')        # for spatial features
library('MODIStsp')  # for downloading NDVI rasters
library('dagitty')   # for directed acyclical graphs
library('ggdag')     # for directed acyclical graphs
library('gratia')    # for ggplot-based GAM figures
theme_set(theme_bw()) # change default theme
source('../functions/betals.r') # betals family written by Simon Wood
source('../analysis/figures/default-figure-styling.R') # for color palettes
source('../earthdata-login-info.R') # personal login info for EarthData
source('../functions/window_hr.R') # function to calculate HRs
```

### 4 Modeling the tapir's movement over time

The script `analysis/tapir/tapirs-moving-window.R` estimates the seven-day home-range size of various tapirs from the Brazilian Cerrado. Here, we simplified the code so that it only estimates the spatial use of the tapir in the manuscript, Anna, which we chose because of the large sample size and high variation in home-range size.

```
# import tapir data from https://github.com/StefanoMezzini/tapirs
anna <- readRDS('../../tapirs/models/tapirs-final.rds') %>%
  filter(name.short == 'ANNA')
anna_tel <- anna$data[[1]] # telemetry data

# re-project using the appropriate UTM projection for the Brazilian Cerrado
ctmm::projection(anna_tel) <- '+proj=utm +zone=22 +datum=NAD83 +units=m'

# calculate the 7-day home-range estimate
window_hr(
  tel = anna_tel,
  window = 7 %#% 'day', # 1 week of data for sufficient sample size
  dt = 1 %#% 'day', # move window over by a single day each time
  fig_path = 'figures',
  rds_path = 'models')
anna_mw <-readRDS('../models/tapirs/CE_31_ANNA-window-7-days-dt-1-days.rds')
anna_mw
```

```
# A tibble: 457 x 13
   date       dataset          guess  model  akde   hr_est_50 hr_lwr_50 hr_upr_50
   <date>     <list>           <list> <list> <list>     <dbl>     <dbl>     <dbl>
 1 2017-06-27 <telemtry[,18]> <ctmm> <ctmm> <UD>       0.555     0.362     0.789
 2 2017-06-28 <telemtry[,18]> <ctmm> <ctmm> <UD>       0.418     0.280     0.583
 3 2017-06-29 <telemtry[,18]> <ctmm> <ctmm> <UD>       0.482     0.337     0.653
 4 2017-06-30 <telemtry[,18]> <ctmm> <ctmm> <UD>       0.597     0.403     0.829
 5 2017-07-01 <telemtry[,18]> <ctmm> <ctmm> <UD>       0.566     0.382     0.786
 6 2017-07-02 <telemtry[,18]> <ctmm> <ctmm> <UD>       0.708     0.459     1.01
 7 2017-07-03 <telemtry[,18]> <ctmm> <ctmm> <UD>       0.642     0.427     0.901
 8 2017-07-04 <telemtry[,18]> <ctmm> <ctmm> <UD>       0.758     0.492     1.08
 9 2017-07-05 <telemtry[,18]> <ctmm> <ctmm> <UD>       0.814     0.534     1.15
10 2017-07-06 <telemtry[,18]> <ctmm> <ctmm> <UD>       0.895     0.574     1.29
# i 447 more rows
# i 5 more variables: hr_est_95 <dbl>, hr_lwr_95 <dbl>, hr_upr_95 <dbl>,
#   t_center <dbl>, posixct <dttm>
```

The `window_hr()` function estimates the tapir's home range using a sliding window approach with a 7-day window (`window = 7 %#% 'day'`) and a one-day slide (`dt = 1 %#% 'day'`). For each set of 7 days, it fits a positional variogram, a continuous-time movement model (Fleming and Calabrese 2021), and a utilization distribution via autocorrelated kernel density estimation (Noonan et al. 2019; Silva et al. 2022). Finally, it saves an exploratory figure (fig. C2) to the `figures` folder and the tibble of times, telemetries, movement models, utilization distributions, and home-range estimates (with 95% confidence intervals) to the `models` folder.

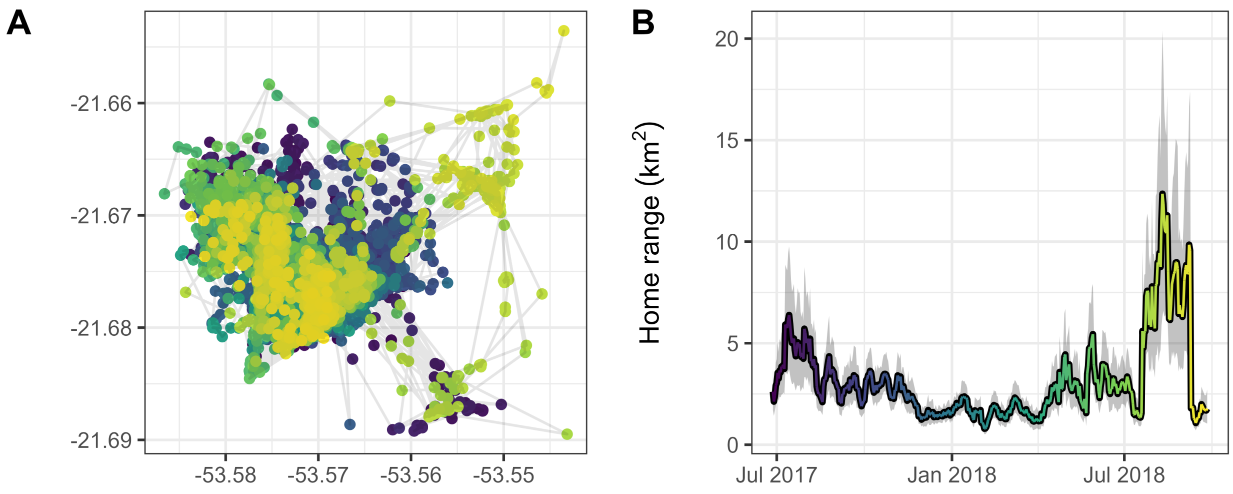


Figure C2: Exploratory figure created by the `window_hr`() function. Panel **A** shows the tapir's GPS locations, while panel **B** shows the seven-day home-range estimates (95% utilization quantile) with 95% confidence intervals.

#### 5 Modeling $\mathbf{E}(R)$ and $\mathbf{Var}(R)$ over time

We estimated the resources in the tapir's habitat using NDVI that we downloaded using the `MODIStsp` package for `R` using the code below.

```
# find the extent of tapir's range
bbox <-
  SpatialPolygonsDataFrame.UD(anna$akde[[1]], # convert to a spatial object
                              level.UD = 0.9995, # utilization quantile
                              level = 0) %>% # no CIs
  st_as_sf() %>%
  st_transform(crs = '+proj=longlat') %>%
  st_bbox()

# download NDVI rasters (if needed, create all necessary folders first)
MODIStsp(gui = FALSE, # do not use the browser GUI, only run in R
         out_folder = 'data/ndvi-rasters/tapir-anna',
         selprod = 'Vegetation Indexes_16Days_250m (M*D13Q1)',
         prod_version = '061', # 2022 raster version
         bandsel = 'NDVI', # NDVI layer only
         sensor = 'Terra', # only terrestrial values, ignore water
         user = USERNAME, # Earthdata username for urs.earthdata.nasa.gov
         password = PASSWORD, # your Earthdata password
         start_date = format(min(anna_tel$timestamp) - 16, '%Y.%m.%d'),
         end_date = format(max(anna_tel$timestamp) + 16, '%Y.%m.%d'),
         spatmeth = 'bbox', # use a bounding box for the extent
         bbox = bbox, # spatial file for raster extent
         out_projsel = 'User Defined', # use specified projection
         output_proj = '+proj=longlat', # download unprojected raster
         resampling = 'bilinear', # raster resampling method for new proj
         delete_hdf = TRUE, # delete HDF files after download is complete
         scale_val = TRUE, # convert from integers to floats within [-1, 1]
         out_format = 'GTiff', # output format
         verbose = TRUE) # print processing messages
```

```r
# save NDVI data as an rds file of a tibble
list.files(path = 'data/ndvi-rasters/tapir-anna/VI_16Days_250m_v61/NDVI/',
           pattern = '.tif', full.names = TRUE) %>%
  rast() %>% # import all rasters as a single stack
  as.data.frame(xy = TRUE) %>% # convert to a data frame
  pivot_longer(-c(x, y)) %>% # change to long format (x, y, name, value)
  transmute(long = x, # rename x column
            lat = y, # rename y column
            date = substr(name, # change name to a date
                          start = nchar('MOD13Q1_NDVI_x'),
                          stop = nchar(name)) %>%
              as.Date(format = '%Y_%j'), # format is year_julian date
            ndvi = value, # rename value column
            dec_date = decimal_date(date)) %>%
  saveRDS('data/ndvi-rasters/tapir-anna/tapir-anna-data.rds')

# import NDVI data
anna_ndvi <-
  readRDS('data/ndvi-rasters/tapir-anna/tapir-anna-data.rds') %>%
  mutate(dec_date = decimal_date(date))
anna_ndvi
```

```
# A tibble: 13,376 x 5
    long   lat date        ndvi dec_date
   <dbl> <dbl> <date>     <dbl>    <dbl>
 1 -53.6 -21.7 2017-06-10 0.626    2017.
 2 -53.6 -21.7 2017-06-26 0.595    2017.
 3 -53.6 -21.7 2017-07-12 0.469    2018.
 4 -53.6 -21.7 2017-07-28 0.421    2018.
 5 -53.6 -21.7 2017-08-13 0.426    2018.
 6 -53.6 -21.7 2017-08-29 0.479    2018.
 7 -53.6 -21.7 2017-09-14 0.440    2018.
 8 -53.6 -21.7 2017-09-30 0.488    2018.
 9 -53.6 -21.7 2017-10-16 0.468    2018.
10 -53.6 -21.7 2017-11-01 0.524    2018.
# i 13,366 more rows
```

We removed the raster for 2017-12-19 because a large portion of the values were unusually low for the region (fig. C3). We hypothesize the change in NDVI was drastic, temporary, and widespread because of a sudden flood. While sudden floods are common for the Cerrado, we believe NDVI was not representative of the available forage availability.

```r
anna_ndvi %>%
  filter(date >= as.Date('2017-08-29'), date <= as.Date('2018-04-07')) %>%
  ggplot() +
  facet_wrap(~ date, nrow = 3) + # a raster for each date
  coord_equal() + # keep the scaling of x and y equal
  geom_tile(aes(long, lat, fill = ndvi)) +
  scale_x_continuous(NULL, breaks = NULL, expand = c(0, 0)) +
  scale_y_continuous(NULL, breaks = NULL, expand = c(0, 0)) +
  scale_fill_gradientn('NDVI', colours = ndvi_pal, limits = c(-1, 1)) +
  theme(legend.position = 'top')
```

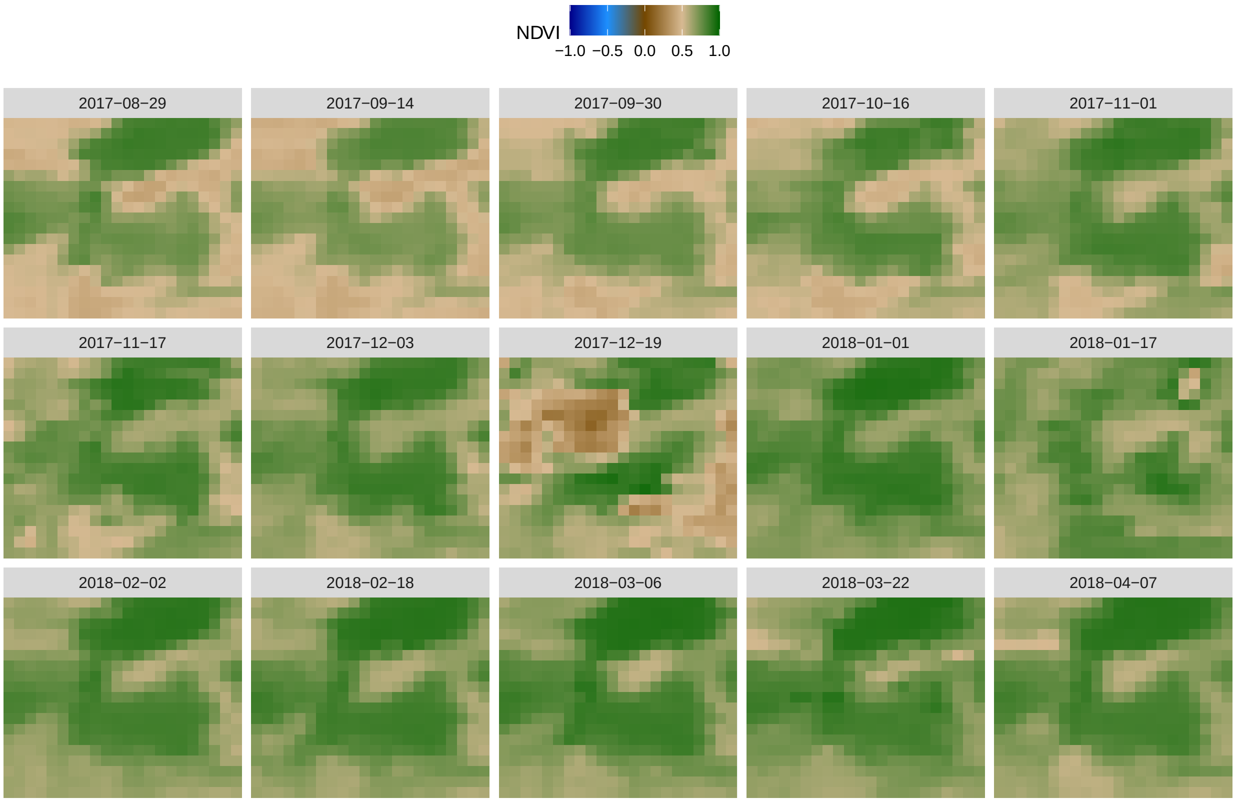


Figure C3: Subset of the NDVI rasters used to estimate the mean and variance in NDVI experienced by the tapir. Notice how many of the values for 2017-12-19 are near zero (brown) but values for the two adjacent rasters are closer to 1 (more green).

```r
anna_ndvi <- filter(anna_ndvi, date != '2017-12-19') # remove biased values
```

Next, we estimate the mean and variance in NDVI using a Generalized Additive Model for location and scale (GAMLS: Stasinopoulos and Rigby 2007) via the `mgcv` package (`family = betals()` in the code chunk below). The `betals` family accepts a list of two predictors: one for the mean parameter, $\mu$, and one for the scale parameter, $\phi$, and it uses logit link functions for both parameters (see fig. C4). The variance of the distribution is a function of both parameters:

$$\sigma^2 = \mu(1 - \mu)\phi. \tag{4}$$

```
m_ndvi <-
  gam(list(
    # mean predictor
    ndvi ~ # not scaling because range is in (0, 1)
      s(long, lat, bs = 'ds', k = 50) + # mean over space
      s(dec_date, bs = 'tp', k = 10), # mean over time
    # scale predictor (sigma2 = mu * (1 - mu) * scale)
    ~
      s(long, lat, bs = 'ds', k = 30) + # scale over space
      s(dec_date, bs = 'tp', k = 10)),  # scale over time
    family = betals(),
    data = anna_ndvi,
    method = 'REML') # REstricted Maximum Likelihood
```

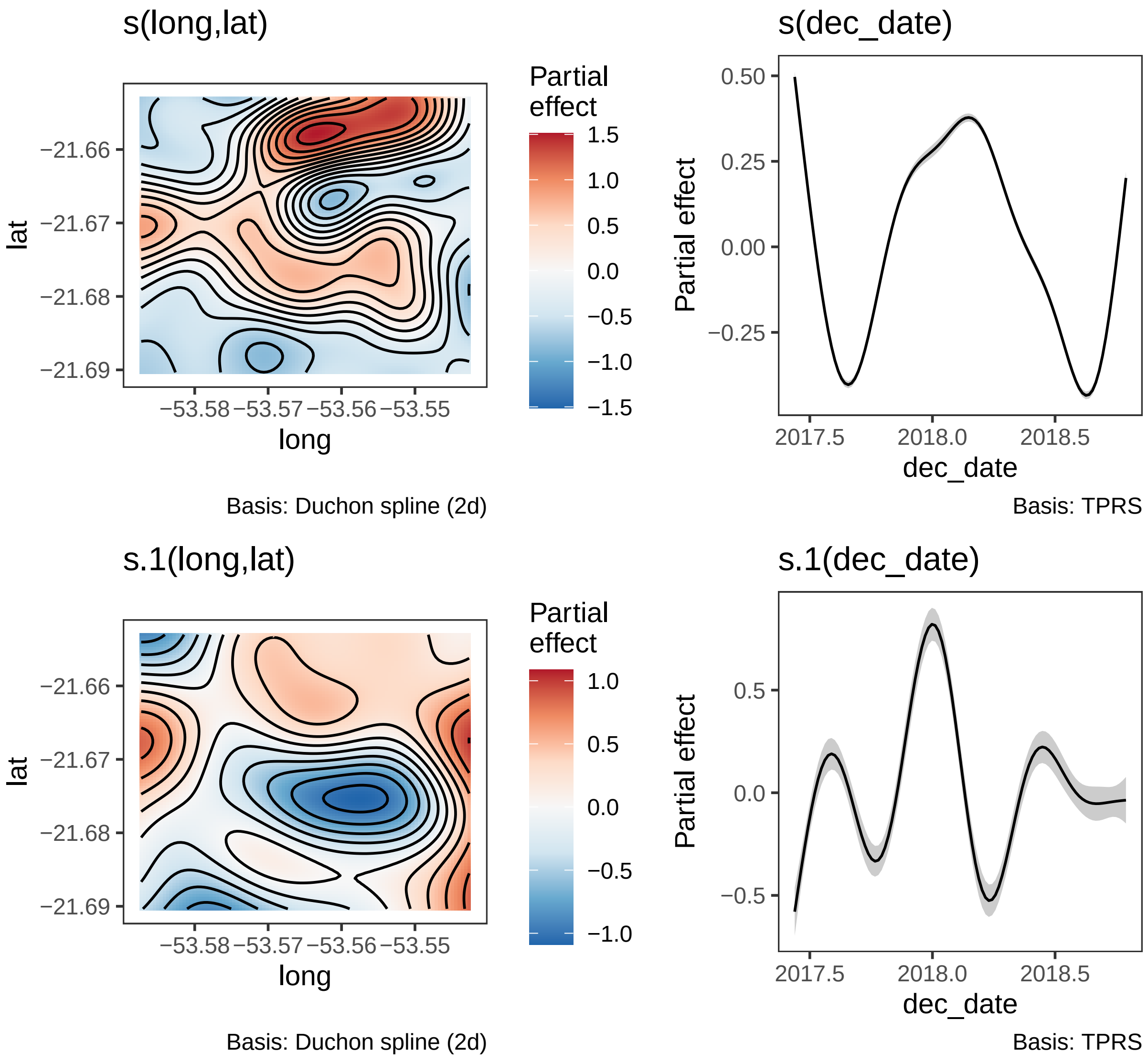


Figure C4: Estimated spatiotemporal trends in mean and scale parameters using the model detailed in the code chunk above. Estimates are provided on the logit link scale. The estimated degrees of freedom for each term can be seen in parentheses in the title of the spatial terms and the y-axis labels of the temporal terms. Shaded ribbons inticate the 95% credible intervals for the temporal terms.

### 6 Modeling the effects of E($R$) and Var($R$) on space use

We start by predicting the mean and variance in NDVI experienced by the tapir at its GPS locations using the beta GAMLS.

```
anna_tel <-
  data.frame(anna_tel) %>% # convert telemetry to data frame
  rename(long = longitude, lat = latitude) %>%
  mutate(dec_date = decimal_date(timestamp)) %>% # needed for predictions
  bind_cols(., # bind telemetry to predictions
            predict(m_ndvi, newdata = ., type = 'response',
                    se.fit = FALSE) %>%
              data.frame() %>% # convert list of predictions to data frame
              # didn't scale NDVI to [0, 1], so no need to back-transform
              transmute(mu = X1, sigma2 = X1 * (1 - X1) * X2)) %>%
  as_tibble()
```

Next, we can estimate the mean and variance in NDVI for each 7-day period using the GPS locations within each period to create the left side of figure 5 from the main manuscript.

```
tapir <-
  readRDS('models/tapirs/CE_31_ANNA-window-7-days-dt-1-days.rds') %>%
  mutate(sub_tel = map(dataset,
                       \(.d) filter(tel, timestamp %in% .d$timestamp)),
         mu = map_dbl(sub_tel, \(.d) mean(.d$mu)),
         sigma2 = map_dbl(sub_tel, \(.d) mean(.d$sigma2))) %>%
  select(date, mu, sigma2, hr_est_95)
```

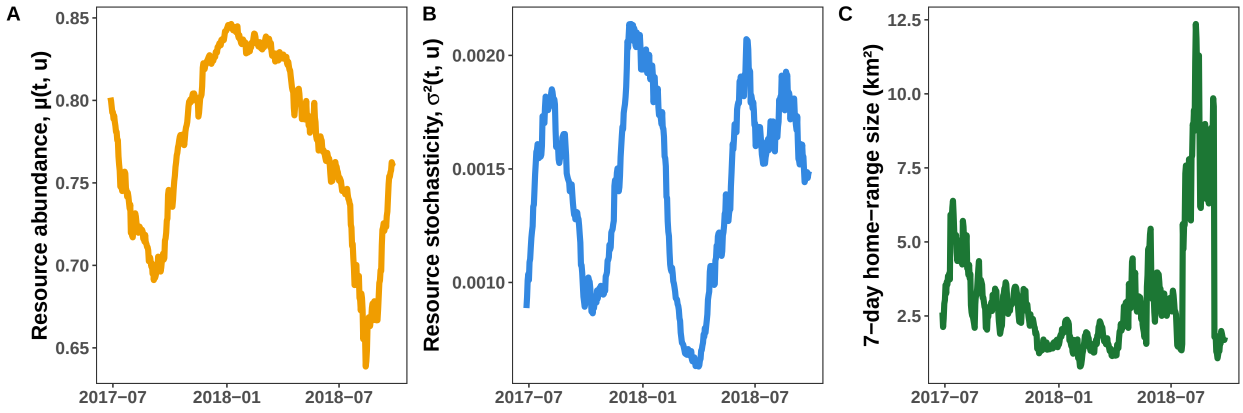

To create the right side of the figure, we need to estimate the effects of E($R$) and Var($R$) on the tapir's space use. To do this, we fit a GAM to the the tapir's 7-day home-range estimates using the mean and variance in NDVI as predictors. As in Appendix B, we provide the causal Directed Acyclical Graph (DAG) in figure C5. See the section on strengths and limitations of the empirical approach in the main text for a discussion of the DAG.

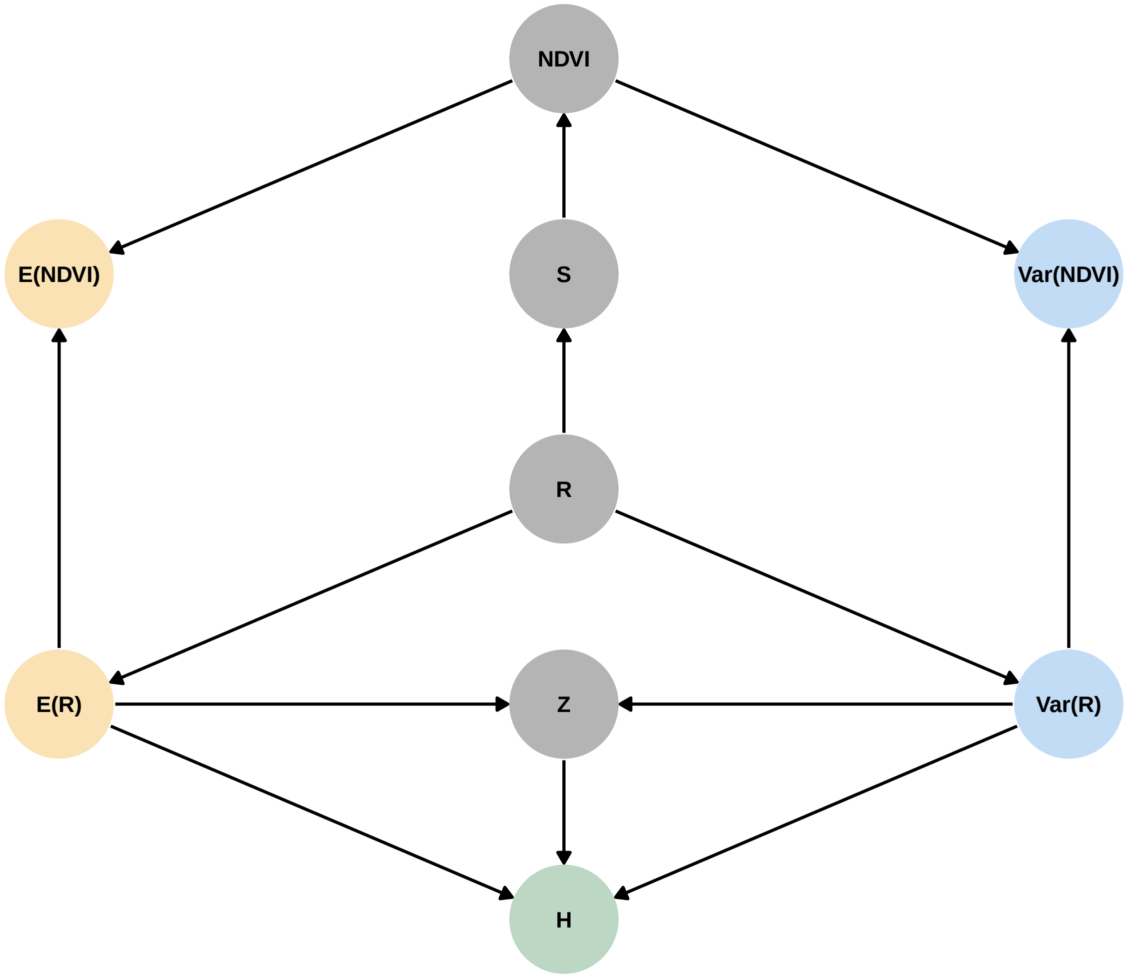


Figure C5: Directed Acyclical Graph assumed for inferring the causal effects of E($R$) and Var($R$) on $H$, where NDVI was used as a proxy for $R$. $Z$ and $S$ indicate unaccounted confounding factors that result from habitat-level variables (e.g., competition, predation, etc.) and satellite-level variables (e.g., noise, cloud cover).

```
m <- gam(hr_est_95 ~ s(mu, k = 4) + s(sigma2, k = 4) + ti(mu, sigma2, k =3),
         family = Gamma('log'), data = tapir, method = 'REML')
draw(m) & theme_bw() + theme(panel.grid = element_blank())
```

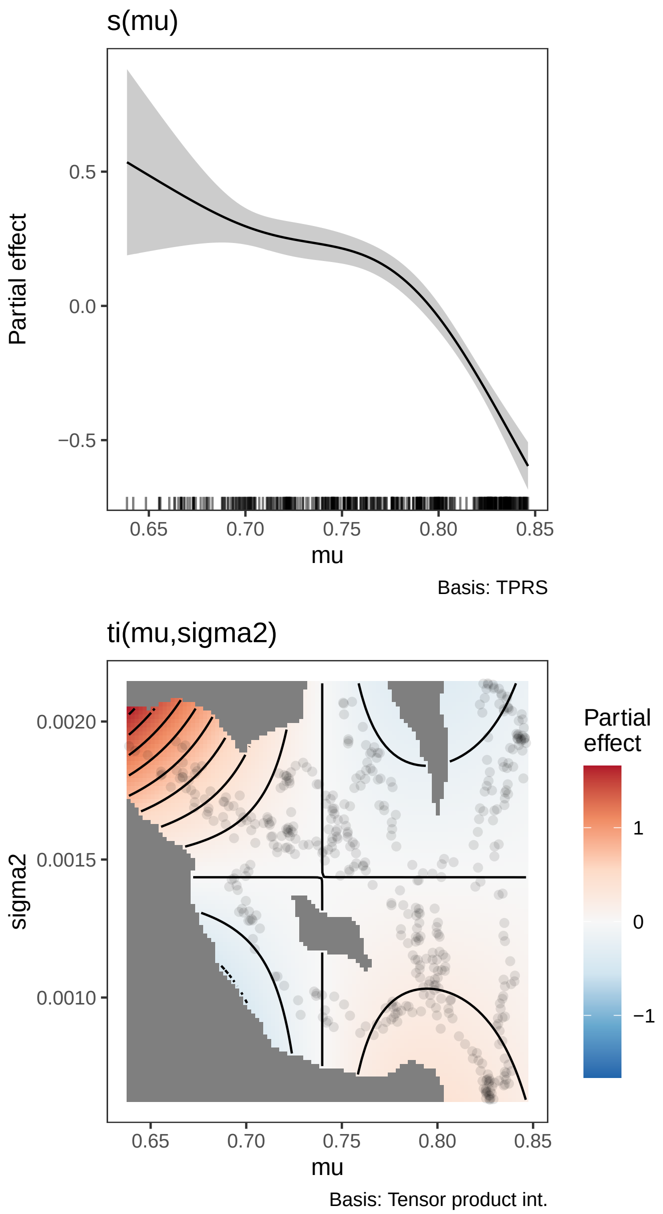


Figure C6: Effects of $\mu(t,u)$ and $\sigma^2(t,u)$ on the tapir's space use (on the log link scale). The estimated degrees of freedom for each term can be seen in parentheses in the y-axis labels. Shaded areas inticate the 95% credible intervals.

We can now predict from the GAM to create the right side of the figure.

```
marginal_preds <-
  tibble(mu = gratia:::seq_min_max(tapir$mu, n = 250),
         sigma2 = gratia:::seq_min_max(tapir$sigma2, n = 250)) %>%
  bind_cols(
    # predictions for the marginal effect of mu
    predict(m, newdata = ., terms = c('(Intercept)', 's(mu)'),
            type = 'link', se.fit = TRUE, unconditional = TRUE) %>%
      as.data.frame() %>%
      transmute(hr_mu_est = exp(fit),
                hr_mu_lwr = exp(fit - 1.96 * se.fit),
                hr_mu_upr = exp(fit + 1.96 * se.fit)),
    # predictions for the marginal effect of sigma2
    predict(m, newdata = ., terms = c('(Intercept)', 's(sigma2)'),
            type = 'link', se.fit = TRUE, unconditional = TRUE) %>%
      as.data.frame() %>%
      transmute(hr_sigma2_est = exp(fit),
                hr_sigma2_lwr = exp(fit - 1.96 * se.fit),
                hr_sigma2_upr = exp(fit + 1.96 * se.fit)))
full_preds <-
  expand_grid(mu = seq(from = floor(min(tapir$mu) * 100) / 100,
                       to = ceiling(max(tapir$mu) * 100) / 100,
                       length.out = 250),
              sigma2 = seq(from = 0.5e-3, to = 2.2e-3, length.out = 250))%>%
  mutate(hr_full_est = predict(m, newdata = ., type = 'response') %>%
           # to avoid excessively large predictions
           if_else(. < 20, ., NA_real_))
```

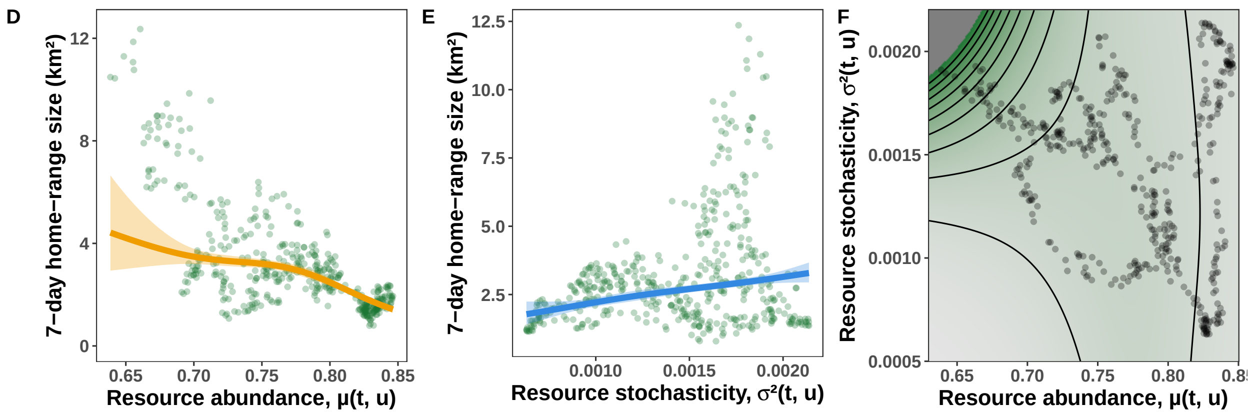
